# Upon microbial challenge human neutrophils undergo rapid changes in nuclear architecture to orchestrate an immediate inflammatory gene program

**DOI:** 10.1101/808568

**Authors:** Matthew Denholtz, Yina Zhu, Zhaoren He, Hanbin Lu, Takeshi Isoda, Simon Döhrmann, Victor Nizet, Cornelis Murre

## Abstract

Differentiating neutrophils undergo large-scale changes in nuclear morphology. How such alterations in structure are established and modulated upon exposure to microbial agents is largely unknown. Here, we found that prior to encounter with bacteria, an armamentarium of inflammatory genes was positioned in a transcriptionally passive environment suppressing premature transcriptional activation. Upon microbial exposure, however, human neutrophils rapidly (<3 hours) repositioned the ensemble of pro-inflammatory genes towards the transcriptionally permissive compartment. We show that the repositioning of genes was closely associated with the swift recruitment of cohesin across the inflammatory enhancer landscape permitting an immediate transcriptional response upon bacterial exposure. These data reveal at the mechanistic level how upon microbial challenge human neutrophils undergo rapid changes in nuclear architecture to orchestrate an immediate inflammatory gene program.

---

The organization of the human genome within the nucleus is central to the control of gene expression and thus cell identity and function. At the largest scale, the genome is folded into chromosome territories that, with the exception of nucleoli, rarely intermingle. However, chromosomes are not randomly distributed across the nucleus. Large and gene poor chromosomes are predominantly positioned at the lamina whereas small and gene rich chromosomes concentrate in the nuclear interior (Fritz et al. 2016). Chromosomes themselves fold into loop domains that physically associate to establish the transcriptionally repressive or inert heterochromatic B compartment or transcriptionally permissive euchromatic A compartment (Dixon et al. 2012; Lieberman-Aiden et al. 2009). The heterochromatic compartment is highly enriched at the nuclear lamina whereas the euchromatic compartment is positioned in the nuclear interior (Kosak et al. 2002).

Loop domains are established in part by the CTCF protein (Dixon et al. 2012; Rao et al. 2014). Convergently oriented pairs of CTCF-bound loci can form CTCF-anchored loops, generated by recruitment of the cohesin complex (Rao et al. 2017; Nora et al. 2017). Cohesin is loaded onto transcribed regions located throughout loop bodies (Busslinger et al. 2017). Once sequestered, the cohesin complex extrudes chromatin in a progressive manner until a pair of convergent CTCF bound sites are reached, a process termed loop extrusion (Fudenberg et al. 2016). Gene promoters connected to transcriptional enhancers by CTCF-mediated loops tend to be highly expressed (Rao et al. 2014), and CTCF occupancy at nearby sites contributes to the maintenance of gene expression and stable chromatin structure (Rao et al. 2017; Nora et al. 2017; Schwarzer et al. 2017; Bintu et al. 2018).

Human neutrophils are abundant, short-lived circulating white blood cells that are critical first-responders to infection and tissue damage. Upon injury or infection, neutrophils exit the circulation via extravasation, migrate towards damaged tissues or infectious foci, phagocytose small pathogens, release reactive oxygen and nitrogen species, and extrude their chromatin as cytotoxic granule-laced extracellular traps (NETs). In addition to their direct role in killing invading pathogens, activated neutrophils rapidly induce the expression of a wide range of cytokines and chemokines to orchestrate an immediate inflammatory response (Ley et al. 2018).

The nuclei of polymorphonuclear (PMN) neutrophils are composed of multiple distinct but internally continuous lobes allowing them to swiftly migrate between (paracellular route) or through (transcellular route) endothelial cells that line blood vessels and interstitial spaces of tissues while maintaining their nuclear integrity (Rowat et al. 2013; Olins et al. 2009; Muller 2013). The Lamin B Receptor (LBR) is an important determinant for imposing a multi-lobular nuclear architecture on neutrophils (Shultz et al. 2003; Hoffmann et al. 2002). Neutrophils of mice deficient in the *Lbr* gene fail to adopt a multi-lobular nuclear shape (Shultz et al. 2003), and mouse neutrophilic cell lines lacking *Lbr* cannot form characteristic toroidal nuclei during differentiation (Zhu et al. 2017). Similarly, humans with *LBR* mutations manifest the Pelger-Huët anomaly, characterized by a reduction in nuclear lobe number in granulocytes (Hoffmann et al. 2002).

Chromatin folding in murine neutrophils is highly enriched for remote genomic interactions, primarily involving heterochromatic regions. These interactions span vast genomic distances resulting in large-scale chromosome condensation. Terminal differentiation of murine neutrophils is also associated with the relocation of centromeres, pericentromeres, telomeres, LINE elements, and ribosomal DNA from the nuclear interior to the nuclear lamina, a process that requires the *Lbr* gene (Zhu et al. 2017). As neutrophils differentiate, the LBR deforms the malleable nuclear envelope by wrapping it around the heterochromatic component of the neutrophil genome, resulting in its characteristic lobed shape.

Upon reaching a tissue site of infection, neutrophils neutralize bacteria in multiple ways: (i) engulfment through phagocytosis; (ii) degranulation to release microbicidal factors into the extracellular space; (iii) release of extracellular traps or NETs that are composed of extruded chromatin fibers and antimicrobial factors; and (iv) rapid induction of cytokine gene expression to coordinate a broader immune response (Ley et al. 2018; Brinkmann et al. 2004). To detect and respond appropriately to diverse invading pathogens, neutrophils express a variety of pattern recognition receptors including cell surface and endolysosomal Toll-like receptors (TLRs), C-type lectin receptors, and formyl peptide receptors, among others. Once activated a variety of downstream signaling pathways converge on the NF-κB and AP1 transcription factors to induce an inflammatory gene program including the cytokines and chemokines IL-8/CXCL8, TNFα, IL-1β, IL-17, and IFNγ (Thomas and Schroder 2013; Garcia-Romo et al. 2011; Tecchio et al. 2014).

The mechanisms by which pathogen sensing pathways interface with the neutrophil genome to induce a rapid and stimulant-appropriate inflammatory gene expression program remain unclear. Here we found that human neutrophil genomes display highly segmented compartments and contracted heterochromatin when compared to human embryonic stem cells. Upon microbe encounter, a specific subset of modestly euchromatic subdomains, spatially segregated from the highly euchromatic A compartment, displayed strengthening of their euchromatic character, and relocalized from a peri-nuclear envelope position towards the nuclear interior. Prominent among the regions that repositioned during human neutrophil activation were gene loci associated with an activated neutrophil-specific gene expression program. Microbial-induced changes in long-range chromatin interactions were globally associated with rapid loss of insulation at euchromatic subdomain boundaries, as well as the formation of *de novo* chromatin loops linking immune response genes to pre-existing and *de novo* formed transcriptional enhancers. The loop-mediated juxtaposition of inflammatory genes to transcriptional enhancers upon microbial exposure was closely associated with the deposition of histone 3 lysine 27 acetylation (H3K27ac), an enhancer-associated histone modification, and rapid loading (<3 hours) of the cohesin complex at the subset of enhancer elements that control an inflammatory gene program. Based on these observations, we propose that the microbe-induced transcriptional signature of activated neutrophils is driven by the re-localization of inflammatory genes to the nuclear interior in conjunction with rapid induction of cohesin-mediated loop extrusion.

## Results

### Human neutrophil development is associated with segmented compartments and contracted genomes

Neutrophil nuclei undergo dramatic morphological changes during differentiation from multipotent progenitors, with terminally differentiated neutrophil nuclei having 3-5 internally continuous but spatially distinct lobes (Supplemental Fig. S1A). To characterize the genomic interactions established during the development of PMN cells, neutrophils were isolated from human peripheral blood, formaldehyde-fixed, and analyzed using *in situ* HiC (Supplemental Table S1) (Rao et al. 2014). The genomes of human neutrophils were slightly enriched for inter-chromosomal interactions when compared to human embryonic stem cells (hESCs) (Fig. 1A). Chromosome territories remained intact and we found no evidence of individual chromosomes being split across multiple lobes (Supplemental Fig. S1B). Notably, compared to hESCs, human neutrophils were depleted for genomic interactions that spanned less than 3Mb but were enriched for interactions that covered more than 3Mb (Fig. 1A).

**Figure 1.**
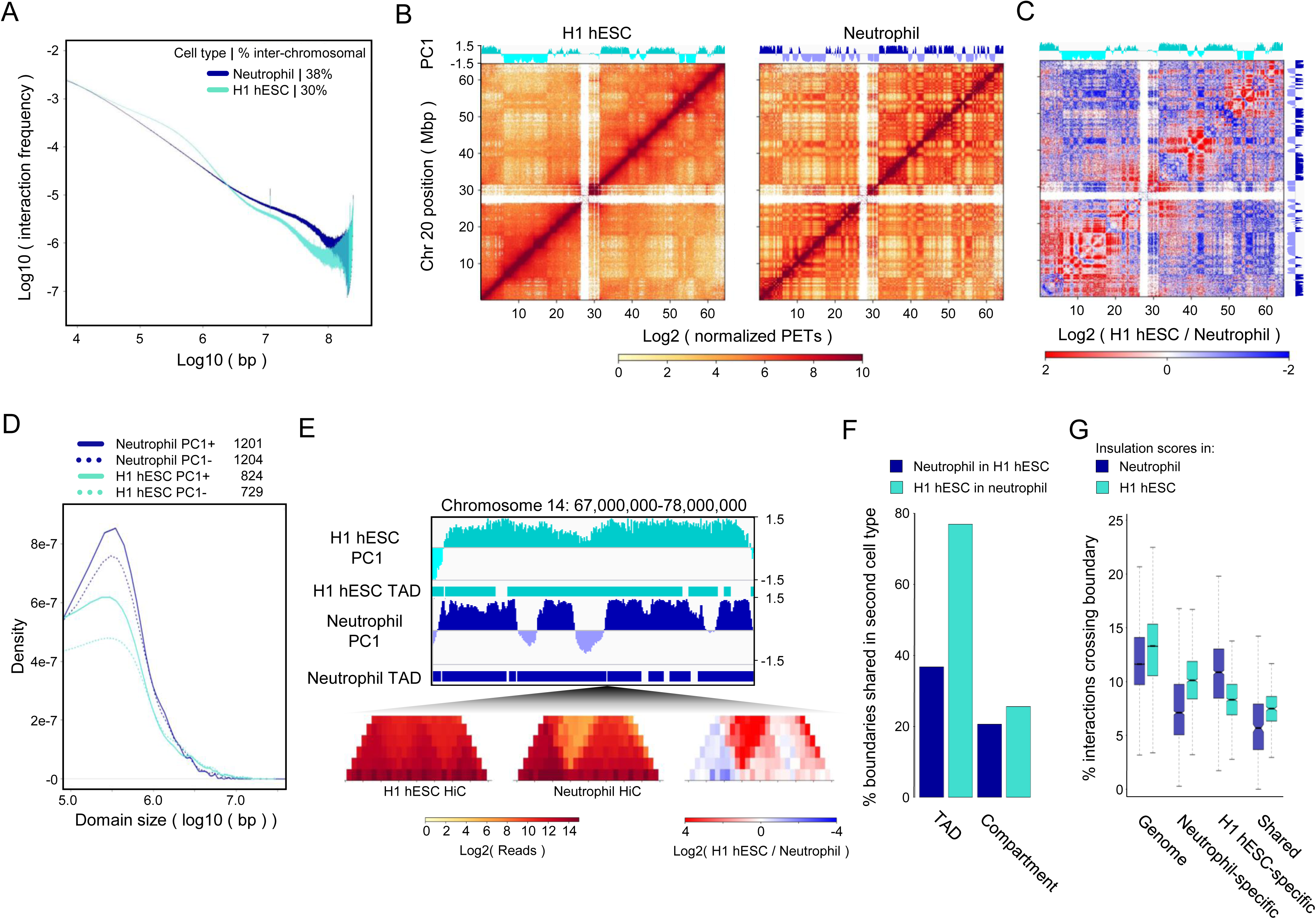
Heterochromatic super-contraction and segmentation of the neutrophil genome during acquisition of the PMN shape. (A) Neutrophil (blue) and H1 hESC (teal) chromatin interaction frequencies as a function of linear genomic distance. Percent interchromosomal paired end tags (PETs) are indicated. (B) Normalized HiC contact matrices for H1 hESC (left) and neutrophil (right) chromosome 20. First principal component 1 eigenvector (PC1) for each HiC matrix is displayed above its respective matrix. Positive PC1 values correspond to the gene-rich A compartment, negative values to the gene poor B compartment, based on human genome build hg38. (C) HiC contact matrix showing the log_2_ fold-change in normalized interactions between the H1 hESC and neutrophil matrices in B, illustrating changes in the organization of chromosome 20 during terminal differentiation from the pluripotent state and acquisition of the PMN shape. PC1 values for H1 hESCs and neutrophils are shown above and to the right of the matrix, respectively. (D) Density plot showing the distribution of A (solid lines) and B (dotted lines) compartment domain sizes in H1 hESCs (teal) and neutrophils (blue). Total number of domains for each data set are listed at top. Domains smaller than 100,000 bp were not considered. (E) Example of a new TAD and compartment boundary formed during differentiation and hyper-compartmentalization of the neutrophil genome. Top (top to bottom): IGV tracks showing H1 hESC PC1 values, H1 hESC TADs, neutrophil PC1 values, and neutrophil TADs. Bottom: Normalized HiC contact matrix of H1 hESC and neutrophil HiC matrices and a log_2_ fold-change difference matrix at a new TAD/compartment boundary on chromosome 14. (F) Percent of TAD and compartment boundaries shared between H1 hESCs and neutrophils. Domain boundaries within 100kb (< 3 windows) were considered shared. (G) Insulation scores in H1 hESCs and neutrophils calculated for each 40kb bin genome-wide, at neutrophil-specific, H1 hESC-specific, and shared PC1 compartment/TAD boundaries for both cell types. Grouped pairs are all significantly different by the Wilcoxon rank sum test with log(p) values <1E-11.

We next constructed contact matrices for hESCs cells and human neutrophils (Fig. 1B). We found that a larger fraction of the neutrophil genome was sequestered in the B compartment when compared to hESCs (Supplemental Fig. S1C). The stereotypic plaid pattern, resulting from the spatial segregation of the A and B compartments, was much more pronounced in human neutrophils compared to hESCs (Fig. 1B). Intra-chromosomal and inter-chromosomal interactions between A and B compartments were both less prevalent in neutrophils versus hESCs (Fig. 1C and Supplemental Fig. S1D). Conversely, long-range genomic interactions across the B compartment were significantly more extensive in human neutrophils than hESCs (Fig. 1C and Supplemental Fig. S1D). During differentiation, large genomic regions that exhibited a continuum of either positive or negative PC1 values in hESCs fragmented into smaller genomic regions that switched PC1 values in neutrophils (Fig. 1C,D). Many of the genomic regions that switched from negative to positive PC1 values during development were associated with a neutrophil-specific transcription signature, whereas those regions switching from positive to negative PC1 values were associated with silencing of lineage-inappropriate genes (Supplemental Fig. S1E-G; Supplemental Table S2). Notably, the hyper-segmentation of compartment domains in the neutrophil genome established *de novo* loop domain and compartment boundaries (Fig. 1E). Specifically, although more than 75% of loop domain boundaries identified in hESCs were conserved in neutrophils, less than 40% of loop domain boundaries in neutrophils were present in hESCs (Fig. 1F). Overall compartment boundaries were poorly conserved between these two cell types (Fig. 1F). Genome-wide analysis of cell type-specific loop domain and compartment boundary element insulation strength confirmed this finding, indicating the existence of cell type-specific boundaries that were associated specifically with either hESCs or human neutrophils, in addition to shared boundaries (Fig. 1G). Taken together our data reveal that human neutrophils, when compared to hESCs, are characterized by a contracted genome with increased enforcement of compartmentalization and highly segmented A and B compartments.

### PMA-induced activation of neutrophils rapidly modulates nuclear architecture

Upon detecting invading microbes, neutrophils rapidly activate an inflammatory-specific transcription signature. As a first approach to examine whether and how the nuclear architecture of neutrophils responds to inflammatory signals, HiC was performed on neutrophils cultured in both the absence and presence of the canonical neutrophil activator phorbol 12-myristate 13-acetate (PMA), a protein kinase C. PMA stimulation of human neutrophils resulted in a global decrease in short-range intra-chromosomal interactions and a global increase inter-chromosomal interactions (Fig. 2A), while exerting minimal effects on A-B compartmentalization and loop domain boundaries (Fig. 2B). Likewise, PMA-induced activation did not trigger large-scale switching of genes or regulatory elements between the A and B compartments (Fig. 2C). However, further scrutiny of chromatin folding across the A compartment revealed a small but significant number of discrete genomic regions that underwent significant PMA-dependent changes from low but positive PC1 values to highly positive PC1 values, indicating an increase in euchromatic character (Fig. 2C). We refer to these regions as PMA ΔPC1 domains (Fig. 2C, Methods). Notably, PMA ΔPC1 domains were strongly enriched for genes implicated in the neutrophil defense response, including genes downstream of key innate immune receptors such as the complement receptors, FCγ receptor, and dectin-2, as well as genes implicated in cell migration and regulation of lysosomal pH (Fig. 2D; Supplemental Table S2).

**Figure 2.**
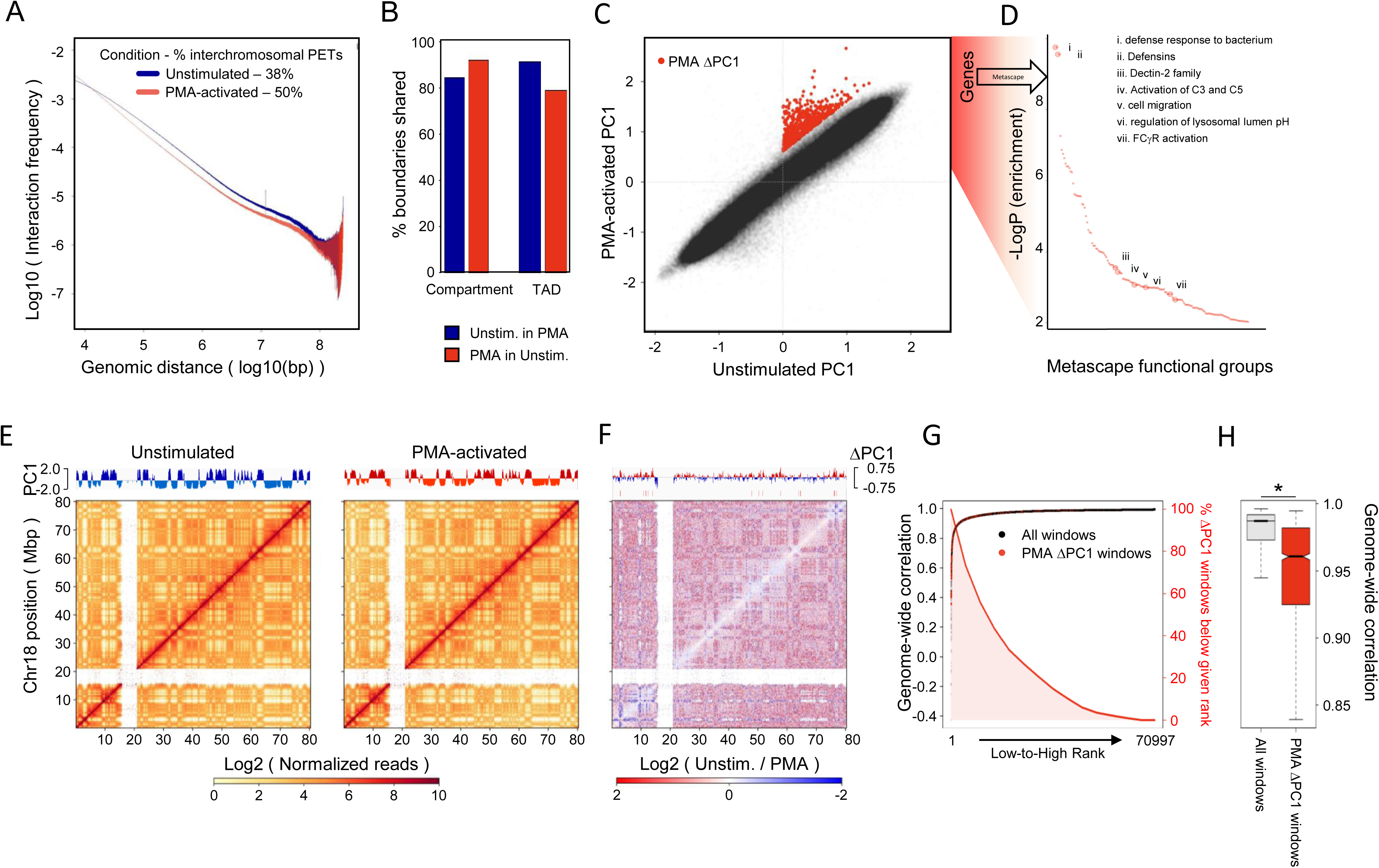
PMA-activation increases euchromatic character at distinct genomic loci encoding for neutrophil activation genes. (A) Unstimulated (blue) and PMA-activated neutrophil (red) chromatin interaction frequencies as a function of linear genomic distance. Percent interchromosomal PETs are indicated. (B) Percentages of TAD and compartment boundaries shared between unstimulated and PMA-activated neutrophils. (C) Scatterplot comparing 10kb-windowed PC1 values between unstimulated and PMA-activated neutrophils. PMA ΔPC1 domains are marked in red. (D) Metascape-defined functional groups enriched for genes found within 100kb of PMA ΔPC1 domains. Complete metascape results can be found in Supplemental Table 2. (E) HiC contact matrices of chromosome 18 for unstimulated (left) and PMA-activated (right) neutrophils, 100kb resolution. Respective PC1 values are shown above each matrix. (F) Difference matrix showing the log_2_ fold-change in normalized interactions between unstimulated and PMA-activated neutrophils. PC1 differences are shown at top, PMA ΔPC1 domains are marked by red ticks. (G) Black and red points: Genome-wide interaction correlations for each 40kb bin in the genome, ranked from most to least differential, left to right. Bins containing PMA ΔPC1 domains are marked with red points. Red line with red shading: Proportion of total PMA ΔPC1 domains found at a given rank or lower, showing a preference for PMA ΔPC1 domains to fall in genomic regions with the most differential chromatin interactions upon PMA stimulation. (H) Boxplots of genome-wide interaction correlation values for PMA ΔPC1 domains and the remainder of the genome during PMA stimulation. Boxplot outliers are not shown. *=Wilcoxon rank-sum test p-value <2E-16.

Analysis of intra-chromosomal HiC contact matrices revealed few significant changes across chromosome 18 following PMA-induced activation (Fig. 2E). PMA ΔPC1 domains, however, often displayed changes in interactions with euchromatin, both in their immediate vicinity, as well as across chromosome 18 (Fig. 2F). Direct measure of genome-wide changes in chromatin organization showed that in activated neutrophils PMA ΔPC1 domains showed large-scale changes in contact frequencies (Fig. 2G). Specifically, ~25% of PMA ΔPC1 domains fell within the top 10% most differentially interacting genomic regions (Fig. 2G). Likewise, PMA ΔPC1 domains on average displayed significantly lower chromatin interaction correlation with unstimulated neutrophils when compared to the genome as a whole (Fig. 2H). Taken together, these data indicate that in PMA-activated neutrophils genic and intergenic domains associated with innate immune genes increase their euchromatic character and undergo alterations in remote genomic interactions.

### Upon microbial exposure a subset of neutrophil inflammatory genes increase their euchromatic character

To validate the alterations in neutrophil euchromatic character using a physiologically relevant stimulus, human neutrophils were cultured in the presence of live *Escherichia coli* bacteria for a period of three hours. *E. coli* co-cultured neutrophils were isolated, formaldehyde cross-linked and analyzed using HiC. Genomes of human neutrophils cultured in the presence of *E. coli* only displayed minor alterations in contact frequencies, maintained overall compartment and loop domain structures (Fig. 3A,B), and remained essentially free of detectable A-B compartment switching (Fig. 3C). However, similar to PMA-activated neutrophils, a distinct subset of genomic regions positioned in the A compartment displayed a substantial increase in euchromatic character upon *E. coli* encounter (Fig. 3C, *E. coli* ΔPC1 domains). Notably, the *E. coli* ΔPC1 domains included genes encoding for cytokines and chemokines, genes associated with neutrophil degranulation, and genes linked with the inflammatory response (Fig. 3D; Supplemental Table S2).

**Figure 3.**
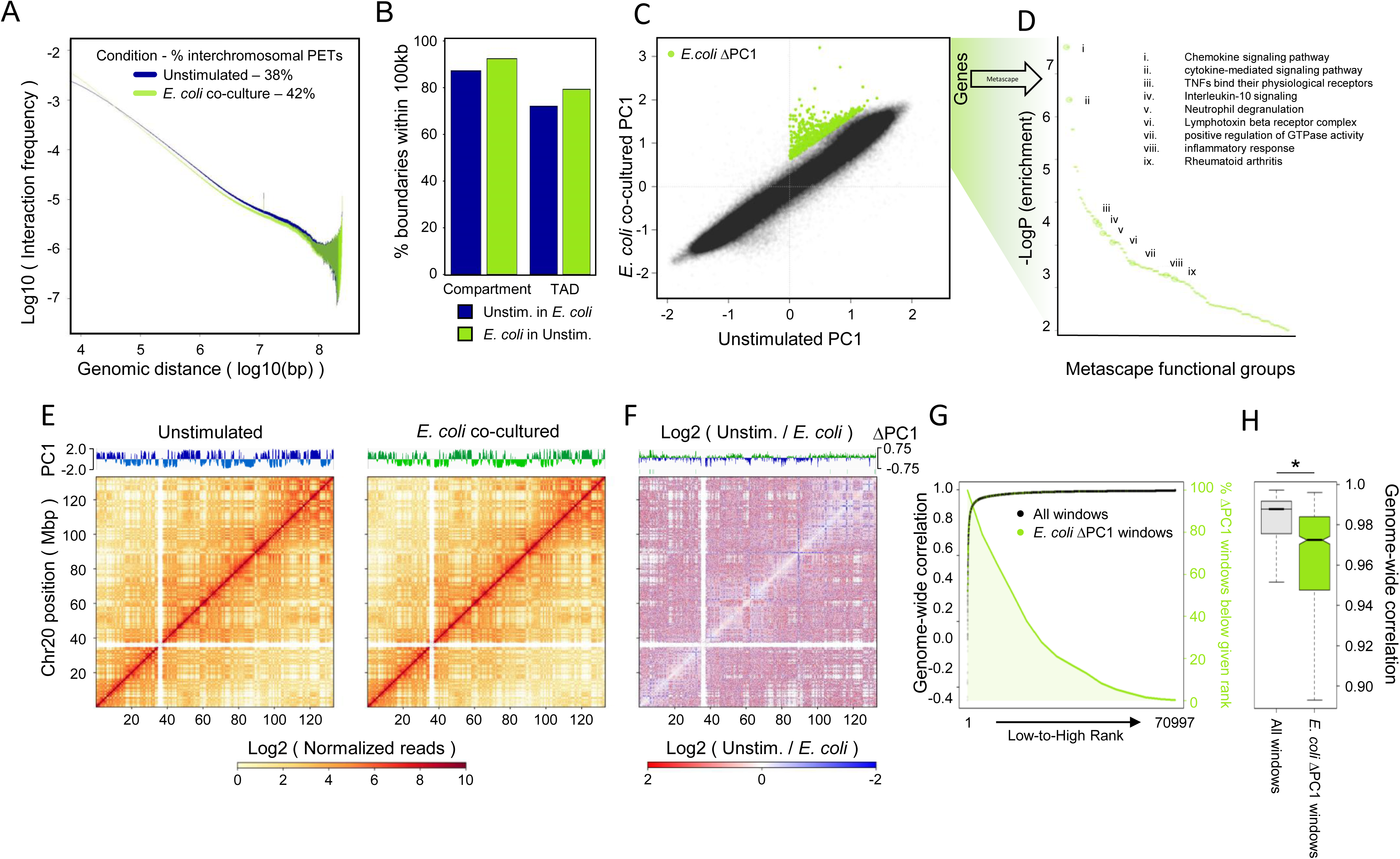
Neutrophil *E. coli* co-culture increases euchromatic character at distinct loci encoding for neutrophil pathogen response genes. (A) Unstimulated (blue) and *E. coli* co-cultured neutrophil (green) chromatin interaction frequencies as a function of linear genomic distance. Percent interchromosomal PETs are indicated. NOTE: Unstimulated neutrophil data is identical to that shown in Figure 2A and is shown here to illustrate differences between unstimulated and *E. coli* co-cultured neutrophil data. (B) Percent of TAD and compartment boundaries shared between unstimulated and *E. coli* co-cultured neutrophils. (C) Scatterplot comparing 10kb-windowed PC1 scores between unstimulated and *E. coli* co-cultured neutrophils. *E. coli* ΔPC1 domains are marked in green. (D) Metascape-defined functional groups enriched for genes found within 100kb of *E. coli* ΔPC1 domains. Complete metascape results are described in Supplemental Table 2. (E) HiC contact matrices of chromosome 20 for unstimulated (left) and *E. coli* co-cultured (right) neutrophils, 100kb resolution. Respective PC1 values are shown above each matrix. (F) Difference matrix showing the log_2_ fold-change in normalized interactions between unstimulated and *E. coli* co-cultured neutrophils. PC1 differences are shown at top with *E. coli* ΔPC1 domains marked with green ticks. (G) Black and green points: Genome-wide interaction correlations for each 40kb bin in the genome, ranked from most to least differential, left to right. Bins containing *E. coli* ΔPC1 domains are marked with green points. Green line with green shading: Proportion of total *E. coli* ΔPC1 domains found at a given rank or lower, showing a preference for *E. coli* ΔPC1 domains to fall within genomic regions with the most differential chromatin interactions upon *E. coli* encounter. (H) Boxplots of genome-wide interaction correlation values for *E. coli* ΔPC1 domains and the remainder of the genome during *E. coli* encounter. Boxplot outliers are not shown. *=Wilcoxon rank-sum test p-value <2E-16.

Similar to PMA-activated neutrophils, *E. coli* co-cultured neutrophils showed few large-scale changes in chromatin organization compared to unstimulated neutrophils (Fig. 3E). *E. coli* ΔPC1 domains, however, showed dramatic increases in genomic interactions involving neighboring euchromatic regions, as well as the remainder of the chromosome upon co-culture with *E. coli* (Fig. 3F). Similar to PMA ΔPC1 domains, *E. coli* ΔPC1 domains were among the most restructured genomic regions in response to *E. coli*, with 25% of *E. coli* ΔPC1 domains assigned to the top 15% of the most differentially interacting regions globally (Fig. 3G). *E. coli* ΔPC1 domains overall displayed significantly lower correlation with unstimulated neutrophil genome structure than the remainder of the genome (Fig. 3H). These data indicate that upon microbial exposure, a subset of genes associated with an inflammatory response increase their euchromatic character.

We next sought to ascertain to what degree ΔPC1 domains differed between stimuli. *E. coli* ΔPC1 domains only partially overlapped with PMA ΔPC1 domains (Supplemental Fig. S2A). The identities of genes in ΔPC1 domains also depended on the stimulus that neutrophils encountered. *E. coli-*specific ΔPC1 domains were highly enriched for chemokine and cytokine genes as well as genes involved in chemotaxis (Supplemental Fig. S2B). In contrast, PMA-specific ΔPC1 regions were enriched for defensin gene clusters (Supplemental Fig. S2B). These data suggest that the changes in euchromatic character regulate stimulant-appropriate inflammatory responses. Supporting this hypothesis, genes residing in stimulus-specific ΔPC1 domains underwent stimulus-specific changes in gene expression. Genes in *E. coli* ΔPC1 domains were more highly expressed upon *E. coli* encounter than upon PMA stimulation, whereas genes in PMA ΔPC1 domains were more highly expressed upon PMA stimulation than during *E. coli* co-culture (Supplemental Fig. S2C). Taken together, these data indicate that neutrophil activation enhances the euchromatic character of a subset of inflammatory response gene loci in a stimulus-dependent manner.

### Rapid assembly and relocalization of a CXCL transcriptional hub upon E. coli encounter

To determine how euchromatic character is strengthened upon microbial activation, we focused on an archetypal *E. coli* ΔPC1 domain containing inflammatory-specific genes encoded within the extended *CXCL* locus. The *CXCL* locus spans a cluster of genes encoding a class of chemokines that include *CXCL8* (IL8), *CXCL1*, and *CXCL2* (MIP2α), each of which is rapidly induced when exposed to microbial agents. We found that in unstimulated neutrophils the *CXCL* locus exists as a loop domain associated with a modestly positive PC1 score which is insulated from neighboring euchromatin (Fig. 4A). Notably, within three hours of exposure to *E. coli*, the euchromatic character of the *CXCL* locus was significantly strengthened (Fig. 4A), accompanied by large scale changes in chromatin folding, with genomic interactions and transcriptional activation spreading into neighboring regions (Fig. 4B,C).

**Figure 4.**
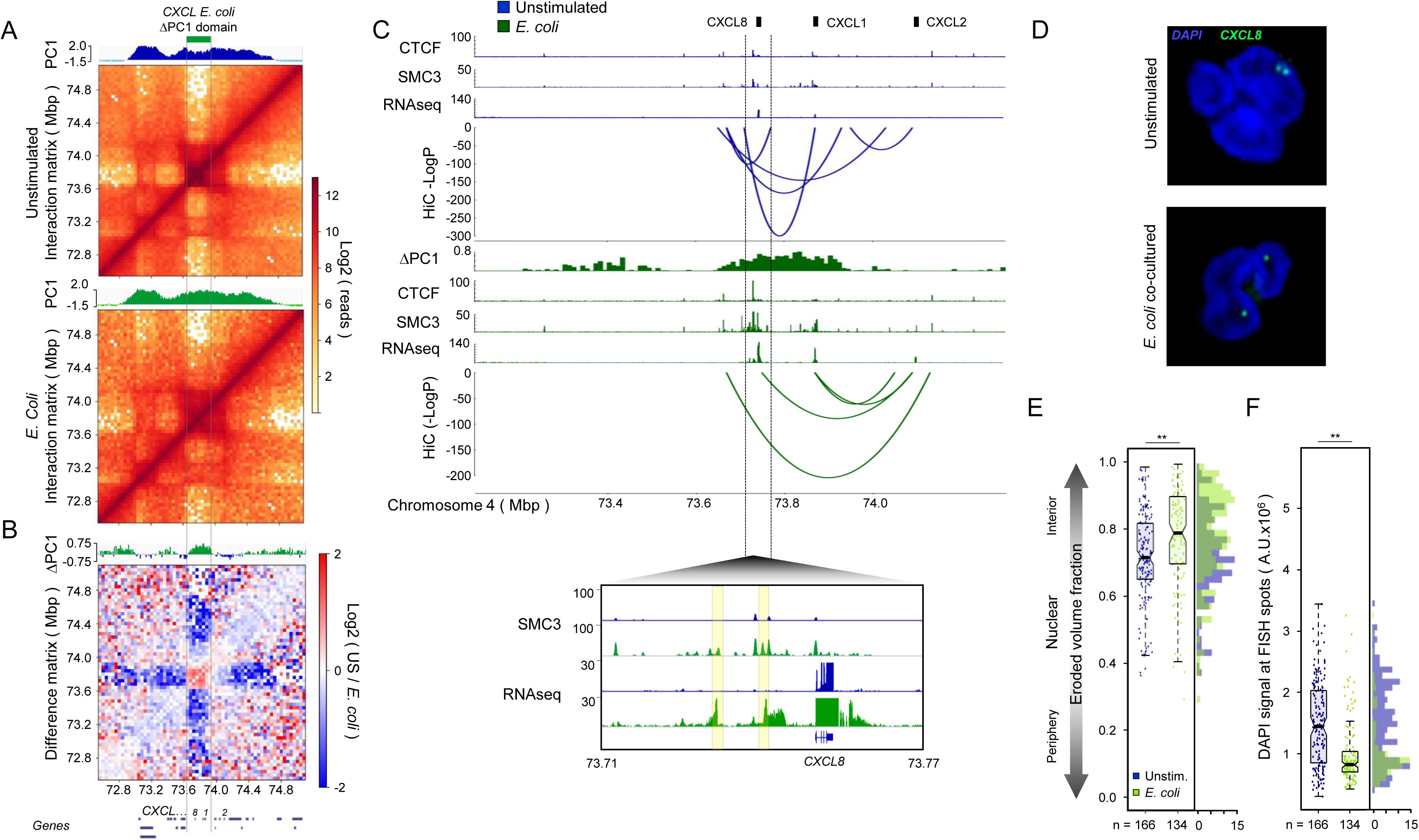
*E. coli* co-culture-induced topological changes at the *CXCL* sub-domain are associated with non-coding transcription, cohesin recruitment, and locus repositioning. (A) HiC contact maps of the extended *CXCL* gene cluster in unstimulated (top) and *E. coli* co-cultured (bottom) human neutrophils. PC1 scores are shown above their respective matrices, *CXCL E. coli* ΔPC1 domain position is noted. (B) Log_2_ difference matrix comparing HiC contacts between unstimulated and *E. coli* co-cultured neutrophils within the extended *CXCL* gene cluster. PC1 differential values (*E. coli* co-cultured – unstimulated PC1 values) are shown above matrix, protein-coding genes in the CXCL gene locus are shown below. (C) Top: Linear genomic features and significant chromatin interactions at the *CXCL E. coli* ΔPC1 domain. CTCF and SMC3 ChIP-seq, RNA-seq, and HOMER-defined chromatin interactions with −log(p) values less than −50 are shown for unstimulated (blue, top) and *E. coli* co-cultured neutrophils (green, bottom) with PC1 differential values (*E. coli* co-cultured – unstimulated PC1 values) shown between. Bottom panel displays SMC3 ChIP-seq and RNA-seq tracks at the *CXCL8* (IL8) gene, demonstrating transcription-associated recruitment of SMC3 to the *CXCL8* locus. Peaks of transcription and associated SMC3 recruitment are highlighted in yellow. (D) Representative FISH image (z-projection) showing the euchromatic *CXCL8* locus (green) in unstimulated (top) and *E. coli* co-cultured (bottom) neutrophils. (E) Quantification of the proportion of nuclear volume between the FISH signal and the nuclear periphery (Eroded volume fraction) in unstimulated and *E. coli* co-cultured neutrophils. Number of alleles analyzed are listed below the boxplots. Wilcoxon rank sum test p-value for the data distributions: ** = <0.00005 (F) Quantification of DAPI signal intensity at the FISH spots identified in B. Wilcoxon rank sum test p-values: ** = <0.00005

To determine whether the alterations in genome folding were associated with gene expression, activated neutrophils were analyzed for transcript abundance as well as CTCF and SMC3 occupancy (Fig. 4C). As expected, CXCL8, CXCL1, and CXCL2 transcript abundance was significantly elevated upon *E. coli* encounter (Fig. 4C). Notably, a recently described non-coding genomic region located immediately upstream of *CXCL8* was also transcriptionally induced upon exposure to bacteria (Fig. 4C) (Fanucchi et al. 2019). While CTCF occupancy was elevated at a site closely linked with the *CXCL8* locus, other CTCF bound sites in the locus were not modulated upon activation (Fig. 4C). In contrast, we found that *E. coli* encounter substantially enriched cohesin occupancy across the locus (Fig. 4C). Cohesin occupancy was particularly prominent at sites closely associated with *de novo* loops that linked the *CXCL8*, *CXCL1*, and *CXCL2* gene bodies, promoter regions, and SMC3-enriched intergenic regions into a shared transcriptional hub (Fig. 4C).

To validate these findings in single cells, we performed fluorescence *in situ* hybridization (FISH) using a probe corresponding to the *E. coli-*specific *CXCL* ΔPC1 domain (Fig. 4D). In unstimulated neutrophils the *CXCL E. coli* ΔPC1 domain localized near the nuclear periphery (Fig. 4D). Upon *E. coli* encounter the *CXCL E. coli* ΔPC1 domain rapidly relocated away from the heterochromatic nuclear periphery towards the nuclear interior, concomitant with its change in euchromatic character and elevated transcript levels (Fig. 4D). Specifically, the *E. coli* ΔPC1 domain relocated from the DAPI-dense portion of the nucleus near the nuclear periphery to the DAPI-sparse nuclear interior (Fig. 4E,F). This change in nuclear positioning was not an indirect result of changes in nuclear morphology, nor activation-induced loss of nuclear integrity, as heterochromatic control probes remained tightly associated with the nuclear periphery during *E. coli* encounter (Supplemental Fig. S3A). Collectively these observations indicate that upon microbial exposure human neutrophils rapidly remodel nuclear architecture to assemble a *CXCL* transcriptional hub in the nuclear interior.

### Neutrophil activation is associated with global loss of insulation at inflammatory genes

The data described above reveal that when human neutrophils encounter bacteria, a subset of inflammatory genes undergo large-scale changes in chromatin folding that spread into neighboring loop domains. To quantitatively describe this loss of subdomain insulation, we computed the insulation scores for genomic regions that surrounded the boundaries of *E. coli* ΔPC1 domains, and upon microbial exposure gained euchromatic character to merge with surrounding euchromatin. We found that upon *E. coli* encounter the gain of euchromatic character across *E. coli* ΔPC1 domains was closely associated with decreased insulation strength at *E. coli* ΔPC1 domain boundaries (Fig. 5A). Although globally the genomic distances separating chromatin interaction anchor points were significantly decreased in activated versus unstimulated neutrophils, the distance separating anchor points of chromatin interactions with *E. coli* ΔPC1 domains increased (Fig. 5B). This loss of insulation at *E. coli* ΔPC1 domain boundaries suggests *de novo* formation of regulatory interactions with the surrounding area (Fig. 5B). Additionally, chromatin interactions contained entirely within *E. coli* ΔPC1 domains were found on average to be significantly stronger in unstimulated neutrophils as compared to *E. coli* co-cultured neutrophils, suggesting a loss of subdomain structure and self-association during microbial encounter (Fig. 5C).

**Figure 5.**
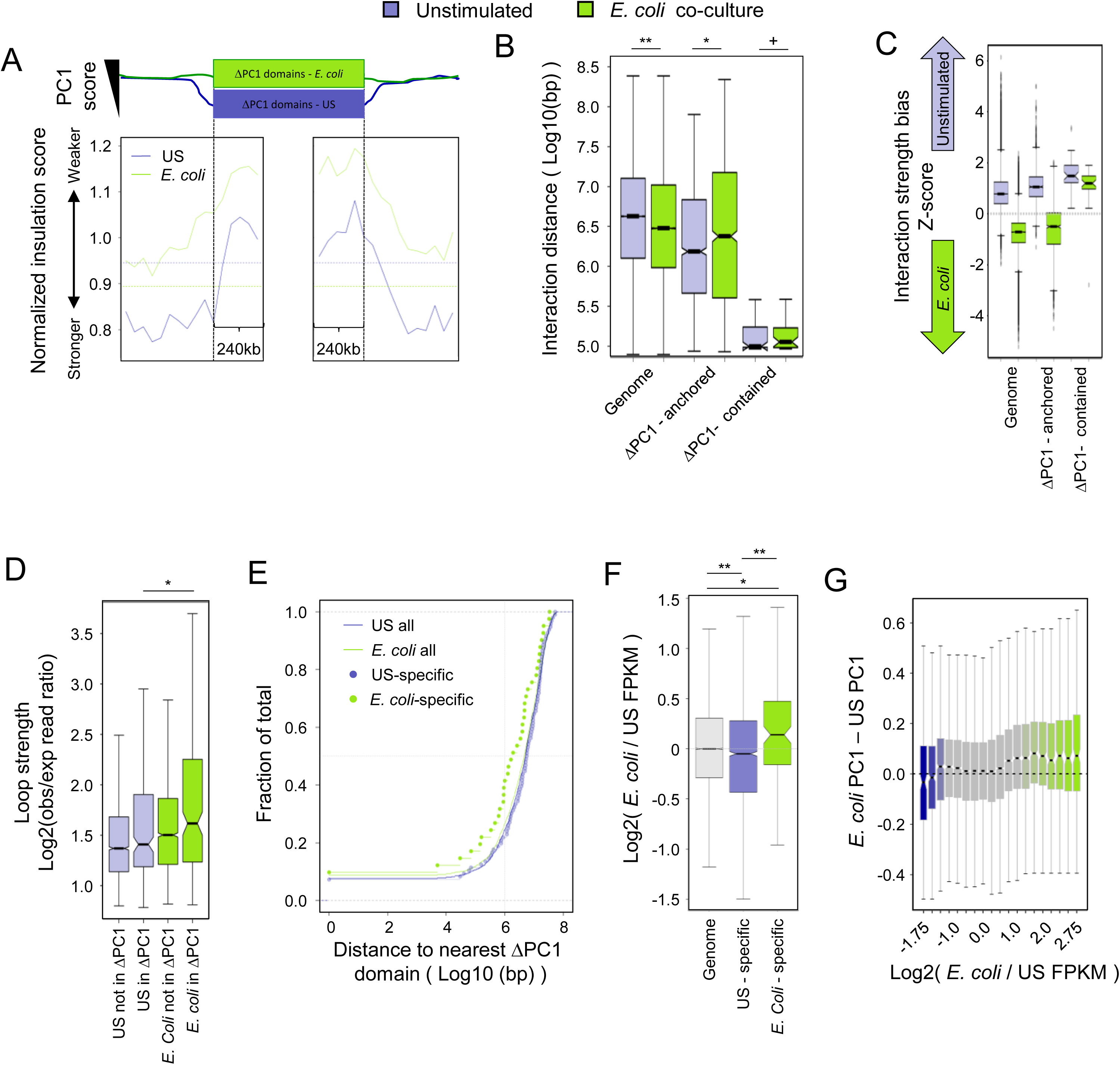
*E. coli* ΔPC1 domains lose local spatial insulation and are associated with increased cohesin-bound chromatin loop strength. (A) Insulation score meta-analysis. Insulation scores were calculated for genomic regions surrounding *E. coli* ΔPC1 domain boundaries. Normalized insulation scores for unstimulated (blue) and *E. coli* co-cultured (green) neutrophils are shown. Vertical dotted lines demarcate meta-domain boundaries. Horizontal dotted lines show median normalized insulation scores at PC domain boundaries genome-wide for each condition. Calculations shown are for all *E. coli* ΔPC1 domains larger than 100kb. (B) Distribution of linear genomic distances between chromatin interaction anchor points genome-wide, for interactions anchored in ΔPC1 domains, and for interactions fully contained within *E. coli* ΔPC1 domains. Wilcoxon rank sum test: **p<2.2E-16; *p<1E-5; +not significant. (C) Interaction strength bias (see methods) of HOMER-defined chromatin interactions for unstimulated and *E. coli* co-cultured neutrophils genome-wide, for those interactions with a single anchor in an *E. coli* ΔPC1 domain, and for those interactions contained entirely within *E. coli* ΔPC1 domains. (D) HICCUPS defined loop strength with respect to *E. coli* ΔPC1 domains in unstimulated and *E. coli* co-cultured neutrophils. (E) Distance between HICCUPS loops and *E. coli* ΔPC1 domains. Differences are not significant by the Kolmogorov-Smirnov test. 5/45 (11%) *E. coli* co-culture specific loops are in *E. coli* ΔPC1 domains, 19/49 are within 1Mb of *E. coli* ΔPC1 domains (45%). *E. coli* ΔPC1 domains make up ~0.3% of the human neutrophil genome. (F) Log_2_ (normalized *E. coli* co-cultured / unstimulated) FPKM values for genes at chromatin interaction anchors shared between unstimulated and *E. coli* co-cultured neutrophils, or at chromatin interactions specific to one condition. Wicoxon rank sum test: **p<1E-4; *p<5E-3. (G) Change in PC1 values at genes with the given mRNA expression differential during *E. coli* encounter.

Given the loss of insulation at *E. coli* ΔPC1 domain boundaries, we next sought to determine the relationship between ΔPC1 domains and gene regulatory chromatin interactions. Although chromatin interactions within *E. coli* ΔPC1 domains were on average weakened during *E. coli* encounter (Fig. 5C), *E. coli* encounter-specific chromatin loops within ΔPC1 domains were significantly stronger than chromatin loops found only in unstimulated neutrophils (Fig. 5D). This finding suggested a gene regulatory role for *E. coli* encounter-dependent loops, and a tight link between these loops and *E. coli* ΔPC1 domains. Supporting this finding, *E. coli*-dependent chromatin loops were generally closer to *E. coli* ΔPC1 domains than were unstimulated neutrophil-specific chromatin loops, and 11% of *E. coli* dependent chromatin loops were identified in *E. coli* ΔPC1 domains, which make up only 0.3% of the genome (Fig. 5E). Importantly, genes near *E. coli* co-culture-specific chromatin loop anchors were significantly more highly expressed than genes at chromatin loop anchors found only in unstimulated neutrophils (Fig. 5F).

Given the enrichment of neutrophil inflammatory response genes in *E. coli* ΔPC1 domains (Fig. 3D) and the link between expression levels and an increase in euchromatic character (Supplemental Fig. S2C), we next determined the relationship between euchromatic character (PC1 score) and transcript levels during microbe encounter. Notably, we found a strong correlation between PC1 score dynamics and transcriptional dynamics, with the most highly induced genes also showing the largest increases in PC1 score, and the most repressed genes showing the largest decreases in PC1 score (Fig. 5G). These phenomena are readily visible at a number of inflammatory loci, wherein the tight self-association of *E. coli* ΔPC1 domains in unstimulated cells is lost in favor of distal regulatory interactions and transcriptional activation during *E. coli* encounter (Supplemental Fig. S4).

Taken together these data indicate that neutrophil transcriptional state, euchromatic character, and spatial localization of genes are closely linked.

### Microbial exposure rapidly recruits cohesin to inflammatory enhancers

Examination of gene regulatory interactions associated with *E. coli* ΔPC1 domains (Fig. 4 and Supplemental Fig. S4) hinted that a large number of *E. coli*-dependent interactions were associated with the recruitment of the cohesin complex to cis regulatory elements. To study this phenomenon and understand its role in *E. coli-*dependent changes in gene expression, we analyzed unstimulated and *E. coli-*exposed neutrophils for SMC3 and CTCF occupancy, as well as changes in H3K27Ac marked enhancer repertoires and transcription. We then focused our analysis on a specific subset of SMC3-amassed enhancers: those H3K27ac-defined enhancers present in *E. coli* co-cultured neutrophils that gained substantial SMC3 occupancy during *E. coli* encounter (Fig. 6A, Methods).

**Figure 6.**
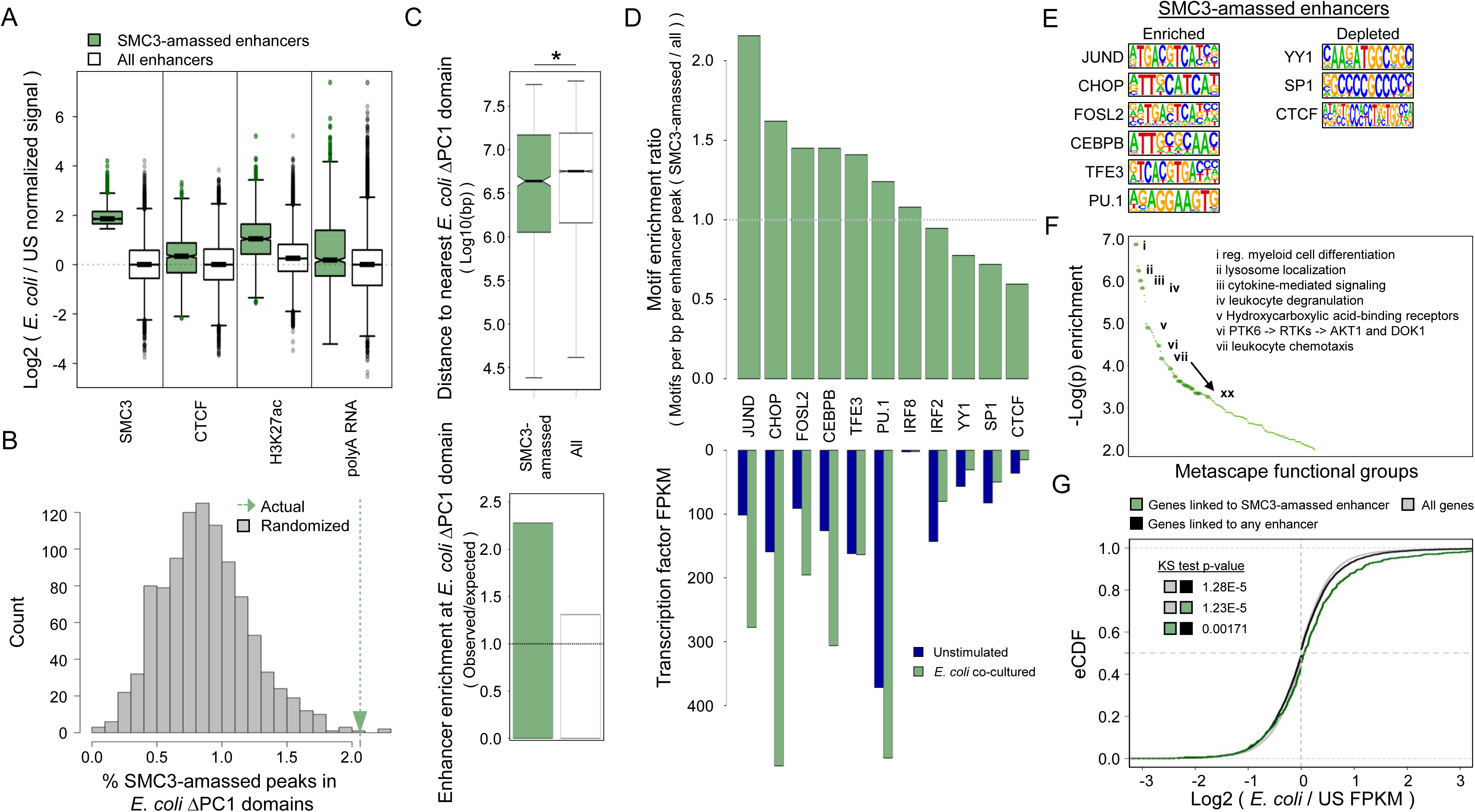
*E. coli* co-culture induces cohesin recruitment to a subset of H3K27ac-defined enhancers. (A) Log_2_ ratio (*E. coli* co-cultured / unstimulated) of the normalized ChIP-seq and RNA-seq signals at H3K27ac-defined enhancers that amass SMC3 as well as all enhancers genome-wide. (B) Histogram showing the percent of SMC3-amassed enhancers falling in *E. coli* ΔPC1 domains randomly positioned within the A compartment (grey), and the percent of SMC3-amassed enhancers falling within actual *E. coli* ΔPC1 domains (green arrow). 997/1000 of random permutations resulted in lower overlap between SMC3-amassed enhancers and ΔPC1 domains than were observed in the empirical data. (C) Top panel shows distance distribution between *E. coli* ΔPC1 domain boundaries and SMC3-amassed enhancers (green) and all enhancers (white). Bottom panel shows observed enrichment of SMC3-amassed enhancers or all enhancers in ΔPC1 domains divided by the expected enrichment of these enhancers in *E. coli* ΔPC1 domains based on 1000 random permutations of *E. coli* ΔPC1 domain positions within the A compartment. * = Wilcoxon rank sum test p-value < 0.005. (D) Top panel indicates ratio of mean transcription factor motif density (motifs per base pair per peak, SMC3-amassed enhancers / all enhancers) for representative transcription factors. Bottom panel shows gene expression values (FPKM) of representative transcription factors in unstimulated and *E. coli* co-cultured neutrophils. (E) Known transcription factor motifs identified in D. (F) Metascape gene functional analysis for genes interacting with SMC3-amassed enhancers. Full metascape analysis results is shown in Supplemental Table 2. (G) Empirical cumulative distribution of log_2_(*E. coli* co-cultured / unstimulated FPKM) values for all genes, genes interacting with any enhancer, and genes interacting with *E. coli* co-culture-dependent SMC3-amassed enhancers.

SMC3-amassed enhancers are characterized by modestly increased CTCF binding, and substantially increased H3K27ac deposition and polyadenylated RNA abundance (Fig. 6A). Supporting the importance of SMC3 and H3K27ac deposition at new regulatory interactions, *E. coli* co-culture-specific chromatin interaction anchors were found to be enriched for SMC3 occupancy and H3K27ac deposition, with only modest changes in CTCF occupancy compared to unstimulated neutrophil-specific interactions (Supplemental Fig. S5A-C). Similarly, H3K27ac-defined enhancers enriched for polyadenylated RNA signal were likewise enriched for SMC3 occupancy and H3K27ac deposition (Supplemental Fig. S5D).

Supporting the importance of SMC3-amassed enhancers in *E. coli* ΔPC1 domain behavior, SMC3-amassed enhancers were enriched at *E. coli* ΔPC1 domains (Fig. 6B). SMC-amassed enhancers were also, on average, localized closer to ΔPC1 domains when compared to the global enhancer repertoire, and were more enriched in ΔPC1 domains than the entire enhancer repertoire (Fig. 6C).

To understand the mechanism of cohesin targeting to SMC3-amassed enhancers, we identified transcription factor binding motifs enriched within both SMC3-amassed enhancers and non-SMC3-amassed enhancers found in *E. coli* co-cultured neutrophils. We then computed the enrichment of transcription factor motif density in SMC3-amassed enhancers compared to all enhancers found in *E. coli* co-cultured neutrophils. Validating this approach, we found that DNA binding motifs associated with known inflammatory regulating factor AP1 including JUND and FOSL2, as well as transcriptional regulators that orchestrate neutrophil differentiation and physiology including CEBPB, CEBP homologue CHOP, and PU.1, were significantly enriched across the bacterial-induced SMC3-amassed enhancer repertoire compared to the entire enhancer repertoire (Fig. 6D, top and Fig. 6E). Notably, transcript abundance associated with these factors was elevated in neutrophils exposed to *E. coli* (Fig. 6D, bottom). Apart from known inflammatory and myeloid regulatory transcription factors, we also found that the bHLH protein TFE3 was enriched at SMC-amassed enhancers when compared to the enhancer repertoire that was not associated with SMC3 occupancy.

Genes interacting with SMC3-amassed enhancers were next analyzed for functional group enrichment. Notably, bacterial-induced SMC3-amassed enhancers were closely associated with genes involved in neutrophil activation, including cytokine signaling and response, chemotaxis, and degranulation (Fig. 6F; Supplemental Fig. S6; and Supplemental Table S2). Analysis of RNA-seq data revealed that genes interacting with enhancers in general showed little preference to be induced upon *E. coli* encounter as compared to any other gene in the genome. In contrast, genes linked to SMC3-amassed enhancers showed a significant increase in gene expression during *E. coli* encounter compared to genes globally, or to genes linked to enhancers in general (Fig. 6G). This phenomenon appears to depend on both SMC3 occupancy and H3K27ac deposition, as interactions with either SMC3-amassed *E. coli*-specific enhancers or SMC3-amassed pre-existing enhancers were both associated with increased gene expression, whereas interactions with enhancers only found in unstimulated cells were not associated with increased gene expression, regardless of SMC3 occupancy (Supplemental Fig. S7).

Taken together, these data indicate that upon bacterial exposure human neutrophils rapidly sequester the cohesin machinery at a specific subset of enhancers to modulate chromatin folding and activate an inflammatory gene transcription program.

## Discussion

The unique morphology of neutrophils has been an enigma since its discovery more than a century ago (Cavaillon 2011). How neutrophil genomes are folded into three-dimensional space and how neutrophil nuclear architecture is altered upon microbial exposure has remained largely unknown. Here we used a genome-wide chromosome conformation capture approach (HiC) to address these questions. We found that human neutrophil nuclei, when compared to embryonic stem cells, displayed a distinct nuclear architecture: (i) a decline in genomic interactions across loop domains (< 3Mb); (ii) a segmentation of large, continuous A and B compartments into numerous small compartments, resulting in the establishment of new compartment and loop domain boundaries; and, (iii) an increase in remote chromosomal interactions across loop domains (> 3Mb). This increase in long-range genomic interactions primarily involved heterochromatic regions indicating a key role for heterochromatic interactions in influencing human neutrophil genome topology. Our data are consistent with previous studies involving murine neutrophils that also displayed a highly contracted genome when compared to progenitor cells and show that key features of neutrophil genome structure are conserved between the murine and human genomes (Zhu et al. 2017).

The neutrophil genome undergoes large-scale alterations in morphology upon bacterial encounter. Using genome-wide chromosome conformation capture studies, we found that such changes involve the repositioning of euchromatic *E. coli* ΔPC1 domains enriched for cytokine and other immune response genes. Upon encountering activating stimuli, these domains gained euchromatic character, repositioning themselves from the nuclear periphery to the more euchromatic nuclear interior. During this process, the boundaries of these domains lost insulation, allowing the domain to merge with neighboring highly euchromatic regions, and further allowing for new chromatin interactions to form and activate an inflammatory gene program. These subdomains resemble a previously identified euchromatic A2 spatial subcompartment positioned between the nuclear periphery and the nuclear interior (Rao et al. 2014; Chen et al. 2018). Based on our observations, we propose that the A2 subcompartment is associated with genes or regulatory elements that need to be transcriptionally repressed, but accessed quickly, precluding both their sequestration to the fully heterochromatic B compartment, as well as their presence in the transcriptionally active A1 compartment.

Our data further provide mechanistic insight as to how neutrophils instruct changes in nuclear positioning and domain insulation upon bacterial encounter. Alterations in chromatin topology both at ΔPC1 domains and across the genome are closely associated with the rapid recruitment of cohesin to a subset of H3K27ac-defined enhancers. While cohesin occupancy is substantially enriched at these enhancers, CTCF binding is only modestly elevated upon bacterial encounter. These observations imply that changes in nuclear architecture are predominantly activated by cohesin-dependent loop extrusion. This finding then raises the question as to how cohesin is being recruited to inflammatory genes upon bacterial encounter. We found that the increase in cohesin occupancy at SMC3-amassed enhancers was closely accompanied by substantial enrichment for the enhancer mark H3K27ac. Hence, we suggest that upon bacterial encounter, human neutrophils activate a signaling response that involves the Toll-like receptor pathway. Motif analysis suggests that Toll-like receptor mediated signaling modulates the expression and/or biochemical activities of key neutrophil-associated transcriptional regulators such as PU.1, CEBP/β, CEBP homolog CHOP, AP1 factors JUN and FOS, as well as TFE3. The activities of such regulators, in turn, would promote the assembly of an active enhancer repertoire as evidenced by the deposition of H3K27Ac, which then rapidly sequesters cohesin at inflammatory response enhancer-gene promoter clusters. Once recruited to SMC3-amassed enhancers, cohesin may act to extrude chromatin until convergent CTCF sites are reached, removing insulation at ΔPC1 domain boundaries by forming *de novo* loop domains in which activated enhancers are placed within close spatial proximity of gene promoters, altogether facilitating the rapid activation of an inflammatory response gene program (Fig. 7).

**Figure 7.**
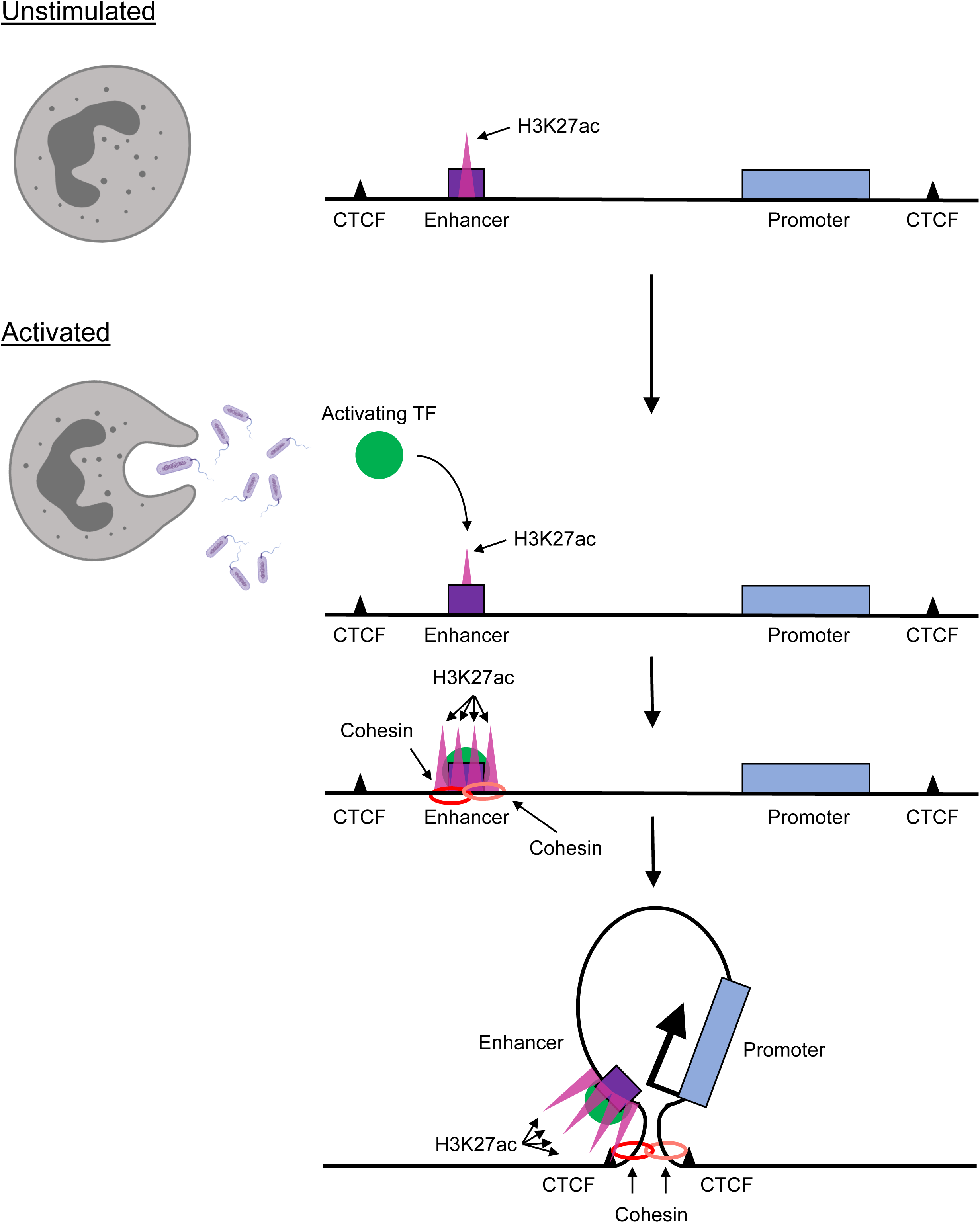
Microbial-induced human neutrophil activation instructs rapid changes in nuclear architecture to orchestrate an inflammatory gene program. Activation-induced transcription factor binding results in H3K27ac deposition, cohesin recruitment, and formation of *de novo* chromatin loops linking enhancers to inflammatory genes to orchestrate an inflammatory gene program.

Why has such an elaborate mechanism of gene activation, including loop extrusion, evolved in human neutrophils? We suggest that segregating enhancers and promoters in spatially distinct loop domains ensures efficient silencing and prevents stochastic activation of an inflammatory specific gene program in unstimulated neutrophils. Only upon exposure to activating stimuli are unstimulated neutrophils instructed to juxtapose the inflammatory enhancer repertoire with their target gene promoters, thus facilitating enhancer-promoter communication and the induction of an inflammatory specific gene program. We hypothesize that the specificity of this response is likely governed by transcription factors downstream of activated receptors that bind their target enhancers, allowing cohesin and histone acetyl transferase recruitment, juxtaposition of target gene promoters, and stabilization of transcription units.

As documented here for human neutrophils during a microbial encounter, enhancers and promoters may be spatially segregated from each other in distinct loop domains until an appropriate environmental signal is received in order to prevent inappropriate or pathological activation of gene expression. Previous studies have documented a related mechanism that orchestrates the developmental progression of lymphoid cells. Specifically, regulatory regions associated with key developmental regulators such as EBF1 and Bcl11b are, in progenitor cells, positioned at the nuclear lamina to suppress premature activation during developmental progression. Upon reaching the appropriate developmental stage, alterations in chromatin folding readily reposition such enhancers away from the transcriptionally repressive environment at the lamina into the euchromatic nuclear interior, leading to assembly of transcriptionally productive enhancer-promoter interactions. The repositioning also directs the enhancer into a single loop domain to facilitate enhancer-promoter communication. Once placed within the euchromatic compartment and within spatial proximity to EBF1 and Bcl11b, enhancers and promoters then act to establish B or T cell identity, respectively (Lin et al. 2012; Isoda et al. 2017). Thus, the inflammatory gene response and activation of a developmental specific gene expression programs share a common mechanism that assures appropriate timing of gene expression.

In sum, here we demonstrate that in human neutrophils, prior to encounter with bacteria, an armamentarium of inflammatory genes was positioned in a transcriptionally passive environment suppressing premature transcriptional activation. Upon microbial exposure, however, human neutrophils rapidly (<3 hours) repositioned the ensemble of pro-inflammatory genes towards the transcriptionally permissive compartment. We found that the repositioning of genes was closely associated with the swift recruitment of cohesin across the inflammatory enhancer landscape permitting an immediate transcriptional response upon bacterial exposure. These data reveal at the mechanistic level how upon microbial challenge human neutrophils undergo rapid changes in nuclear architecture to orchestrate an immediate inflammatory gene program.

## Materials and methods

### Human subject details

Blood for neutrophil isolation was obtained via venopuncture from healthy human volunteers under written informed consent approved by the UC San Diego Human Research Protection Program (#131002X).

### Blood draws and neutrophil isolation

Whole blood was layered onto Polymorphprep reagent (Accurate Chemical and Scientific Corp., AN1114683) and was centrifuged for 45 minutes at 500g, 25°C, and allowed to stop without braking. The granulocyte layer was extracted and contaminating red blood cells were lysed as needed (generally 1-3 times) with brief resuspensions in sterile H2O followed by immediate flooding with 1x phosphate buffered saline (PBS) and centrifugation at 500g for 7 minutes at 25°C. Cells were checked for purity via Wright-Giemsa staining; the final granulocyte fraction was generally >95% neutrophils. For RNA-sequencing experiments, neutrophils were further purified to homogeneity using an EasySep Human Neutrophil Enrichment kit (Stemcell technologies, 19257) as per the manufacturer’s protocol.

For all experiments neutrophils were cultured in HBSS +Ca/+Mg/-Phenol red (Thermo Fisher, 14025092) with the addition of 0.5% endotoxin-free BSA (Akron, AK8917-0100) at 37°C in a 5% CO2 humidified incubator.

### Wright-Giemsa staining

Neutrophils (1×10^5) were spun onto cover slips using a Cytospin3 (Shandon, 74010121 GB) and flooded with Wright stain (Sigma, WS16-500ML) for 3 minutes. Cover slips were then washed with six consecutive dips in water baths. Cover slips were then allowed to air dry and were then flooded with Giemsa stain (Sigma, GS500-500ML) and allowed to incubate for 7 minutes before being washed as above and allowed to air dry.

### Neutrophil activation

Neutrophils were plated at the desired cell numbers and treated with 25nM phorbol 12-myristate 13-acetate (PMA, Promega V1171) or co-cultured in the presence of *E. coli* strain K1 at a multiplicity of infection (MOI) of 5. Stimulations were performed for 3 hours and cells were harvested as detailed below.

### Chromatin immunoprecipitation with sequencing (ChIP-seq)

Neutrophils were plated at 10−20×10^6^ cells/10mL for each ChIP experiment. At the completion of each experiment cells were washed with fresh media, formaldehyde was added to the culture to a final concentration of 1%, and cells were cross linked with agitation for 10 minutes at room temperature. Fixation was then quenched for 5 minutes with glycine at a final concentration of 0.13M. Fixed cells were scraped from the plate and washed three times in ice cold 1x phosphate buffered saline (PBS) with 0.1mM EDTA and 1x EDTA-free complete protease inhibitors (Roche 05056489001). Cell pellets were snap frozen in liquid nitrogen and stored at −80°C until processing.

To bind antibody to ProteinG Dynabeads (Invitrogen 10004D), beads were washed 3×1mL with bead wash buffer (1x PBS, 5mg/mL BSA, Roche complete EDTA-free protease inhibitor, 0.22uM filtered) and resuspended in 500uL of the same. 1-5ug of antibody was added and allowed to bind beads overnight, rotating at 4°C. The following day beads were washed 3×1mL with bead wash buffer and resuspended in 100uL RIPA 150 (50mM Tris pH 8.0, 150mM NaCl, 0.1% SDS, 0.1% Sodium deoxycholate, 1% TritonX100, 1mM EDTA).

For each ChIP, cells were thawed and lysed on ice for 10 minutes with inversion in Farnham lysis buffer (5mM PIPES pH 8.0, 85mM KCl, 0.5% NP-40, 10mM EDTA, protease inhibitors) with or without 20 draws through an 18 gauge needle. Nuclei were spun down for 5 minutes, 2000rpm at 10°C in a bench top microfuge, supernatant was discarded, and nuclei were resuspended in 300uL RIPA 150. Chromatin was then sonicated in a Diagenode Bioruptor 300 chilled to 4°C for 3×8 cycles of 30” on and 30” off, set on high with 5 minutes of cooling time between each set of 8 cycles. The insoluble fraction was spun down for 20 minutes at maximum speed, 4°C, in a benchtop microfuge. Input and IP samples were split to separate new tubes, IP volume was adjusted to 900uL with RIPA 150, and 100uL of Protein G dynabeads bound to the antibody of interest in RIPA 150 was added to each IP. Chromatin was allowed to bind to antibody-bead conjugates overnight while rotating at 4°C. Following binding, beads were washed 2×5 minutes in RIPA 150, 2×5 minutes in RIPA 500 (50mM Tris pH 8.0, 500mM NaCl, 0.1% SDS, 01% Sodium deoxycholate, 1% TritonX100, 1mM EDTA), 2×3 minutes in LiCl wash (10mM Tris pH 8.0, 250mM LiCl, 1% NP-40, 1% Sodium deoxycholate, 1mM EDTA), and once in 1x TE. Beads were transferred to clean tubes at the start of each new wash buffer. DNA was eluted from beads with 200uL elution buffer (1mM Sodium carbonate, 1% SDS) for 1 hour, 65°C with shaking, at which point beads were removed and cross-links were reversed overnight at 65°C. Eluted DNA was purified using a ChIP DNA clean and concentrator kit (Zymo D5205).

DNA for ChIP and other high throughput sequencing approaches was processed as follows: End repair was performed using an Epicenter End-It kit (Lucigen ER0720), according to manufacturer’s instructions and column purified in a Zymo Minelute column (Zymo D4013). A-tails were added by incubating DNA in 1x NEB buffer 2 (New England Biolabs B7002S) with the addition of 200uM dATP and 7.5 units Klenow (exo-) (New England Biolabs M0212L) at 37 degrees for 45 minutes. NEB Next adaptors (New England Biolabs E7337A) were ligated using an NEB quick ligation kit (New England Biolabs M2200L) at benchtop temperature for 30 minutes followed by treatment with 2uL USER enzyme (New England Biolabs M5505L) for 15 minutes at 37°C. DNA was purified using an AmpureXP bead-analogous two-step SPRI bead protocol (Rohland and Reich 2012), resulting in purification of DNA fragments between ~200-800 bp.

PCR amplification of final libraries for sequencing was performed with Phusion hot start polymerase II system (ThermoFisher F549L) in conjunction with the NEB Next indexing system (New England Biolabs E7335L and E7500S). Final size selection for all high-throughput sequencing libraries was performed using a home-made two-step SPRI bead-based DNA purification system, resulting in final DNA fragment sizes of ~200-800bp.

### RNA-sequencing

At specified time points neutrophils were washed once with PBS and lysed in the RLT buffer component of the Qiagen RNeasy mini kit (Qiagen 74106) with the addition of 10ul/mL 2-mercaptoethanol, homogenized via Qiashredder (Qiagen 79654), and snap frozen in liquid nitrogen. Total RNA was purified via RNeasy mini kit (Qiagen 74106) according to the manufactures instructions, including the RNase-free DNase (Qiagen 79254) treatment step. RNA was eluted in H2O, Turbo DNase kit buffer (ThermoFisher/Ambion AM1907) was added to a 1x concentration and RNA was treated with 4 units of Turbo DNase at 37degrees for 30 minutes. Turbo DNase was then treated with inactivation reagent per manufacturer’s specifications. mRNA was purified from total RNA using a Dynabead mRNA purification kit (Life Technologies 61006). First strand synthesis was performed using SuperScript III first strand synthesis system (ThermoFisher 18080051) as follows: 100-500ng RNA; 0.5uL Oligo(dT) primer; 0.8uL random hexamer; 1uL 10mM dNTP, H2O to 9.5uL. Mixture was incubated to 70 degrees for 10 minutes then snap frozen. First strand synthesis mix composed of 2uL 10x RT buffer, 4uL 25mM MgCl2, 2uL 0.1M DTT, 0.5 uL of 120ng/uL ActinomycinD, 40U RNaseOUT, 200U SuperScriptIII was added to the mixture which was then incubated at 25 degrees for 10 minutes, 42 degrees for 45 minutes, 50 degrees for 25 minutes, and 75 degrees for 15 minutes. Unincorporated nucleotides were removed from the mixture using a ProbeQuant G-50 column (Sigma GE28-9034-08). First strand synthesis reaction as then brought to 51uL with H2O and cooled on ice. 24uL of second strand mixture composed of 1uL 10x RT buffer, 2uL 25mM MgCl2, 1uL 0.1M DTT, 2uL of 10mM dATP, dGTP, dCTP, dUTP mix, 15uL of 5x second strand synthesis buffer (New England Biolabs B6117S), 0.5uL E. coli ligase (NEB M0205S), 2uL DNA polymerase I (NEB M0209S), and 0.5uL RNaseH was added and mixture was incubated at 16 degrees for 2 hours. DNA was purified using a DNA clean and concentrator kit (Zymo D4013) and sonicated on a Covaris E220 with the following settings; Duty cycle 10%; Intensity 5; Cycle per burst 200; Time (seconds) 180. Sonicated DNA was purified using a DNA clean and concentrator kit. DNA was prepared for high throughput sequencing using the methodology described above for ChIP-seq, with the addition of 1uL UNG (ThermoFisher/Applied Biosystems N8080096) during USER enzyme treatment.

### Whole genome bisulfite sequencing

Neutrophils were washed twice with PBS and genomic DNA was isolated using a DNeasy Blood and Tissue kit (Qiagen 69504). 1 μg of genomic DNA mixed with unmethylated lamda DNA at a concentration of 0.5% of total DNA was sonicated by Biorupter 300 with 20 cycles (30 seconds on and 30 seconds off at low power). Fragmented DNA was end-repaired and A-tailed as described above. TruSeq adapters (Illumina FC-121-2001) were ligated to fragmented DNA which was then purified by running on a 2% agarose gel. Bisulfite conversion was performed using the MethylCode kit as described by the manufacturer (Invitrogen MECOV-50). Bisulfite-treated DNA was amplified by using a TruSeq PCR primer mixture and Pfu Turbo Cx Polymerase, agarose gel purified, and sequenced on an Illumina HiSeq 2500 sequencer with paired end 150bp reads.

### E. coli culture and MOI determination

*E. coli* strain K1 was grown in LB at 37 degrees with shaking overnight, and diluted into a fresh culture and grown to exponential phase the day of each experiment. *E. coli* was then pelleted at 3000rpm for 10 minutes at 10 degrees on a benchtop centrifuge, washed in cell culture media, and added to neutrophil cultures at an MOI of ~5 in HBSS +Ca/+Mg/-Phenol red with 0.5% endotoxin free BSA. 9 1:10 serial dilutions of *E. coli* containing media were plated on LB agar and grown overnight at 37 degrees. The resulting colonies were counted in order to assess MOI for individual experiments.

### In situ HiC

*In situ* HiC was performed as described (Rao et al. 2014), modifying only the MboI restriction enzyme digest time to assure proper digestion of chromatin. Generally, HiC libraries prepared from activated neutrophils were digested for 2-4 hours with 50 units MboI to avoid over-digesting the chromatin. The remainder of the library preparation adhered to the published protocol and reagents exactly. HiC library DNA was prepared for high throughput sequencing using the NEB Next platform according to manufacturer’s instructions, and sequenced using paired end 100bp reads.

### Fluorescence in situ hybridization (FISH)

Cover slips were incubated overnight in 1%HCl in 70% ethanol, washed 3x with H2O, once in 70% ethanol, and stored in 100% ethanol. Coverslips were allowed to air dry prior to adding cells. Cells were incubated on cover slips in 24 well plates as described above. At the completion of incubation times, cells were washed 3×3 minute in PBS and fixed for 30 minutes in 6% paraformaldehyde (Electron Microscopy Sciences 15710) in 1x PBS. PFA was flushed out with >5 volumes of PBS/0.05% Tween-20 (PBST), ensuring that cells never contact the air. Residual PFA was quenched via incubation with fresh 20mM glycine in PBS for 15 minutes at room temperature. Cells were permeabilized in PBS+0.5% Triton x-100 for 20 minutes at room temperature, washed twice with PBST, and incubated in PBS + 100ug/mL RNase A (Qiagen 19101) for 1 hour at 37°C. Cells were then treated with 0.1N HCl for 5 minutes at room temperature, washed 2×3 minutes with 1xPBS, 2×5 minutes with 2xSSC, then incubated for >48 hours in 2xSSC/50% formamide at 4°C. Cover slips were then blotted dry and 5uL probe containing 75-200ng labeled DNA was added to each coverslip. Cover slips were then sealed on top of glass slides along with probe using rubber cement. Probes and genomic DNA were denatured together for 5 minutes at 78°C on a heat block and allowed to hybridize for 16-48 hours at 37°C. Following hybridization cover slips were washed 1×15 minutes in SSC/50% formamide pre-warmed to 37°C, 3×15 minutes in 2xSSC pre-warmed to 37°C, 3×7 minutes in 0.1xSSC pre-warmed to 60°C, 3×7 minutes in 4xSSC/0.02% Tween-20 pre-warmed to 42°C, 1×5 minutes with 2xSSC pre-warmed to 37°C, and 2×5 minutes in 1x PBS. Cells were then post-fixed in 4% PFA in 1x PBS for 10 minutes at room temperature, and PFA was flushed out as above. Cells were washed 1×10 minutes in PBST+DAPI, 4×5 minutes 1xPBS, and mounted in Prolong Gold mounting media (ThermoFisher P36930).

FISH probes were prepared from bacterial artificial chromosomes (BACs) using nick/translation (Roche 11745808910). 1ug BAC DNA was used in each 20uL nick/translation reaction along with the following fluorophores, as needed: ChromaTide Alexafluor 488-5-dUTP (ThermoFisher/Life Technologies C11397), Cy3-dUTP (VRW 42501), or AlexaFluor 647-aha-dUTP (ThermoFisher/Life Technologies A32763). Nick/Translation was performed for 5-16 hours at 15 degrees and terminated by addition of 1uL 0.5M EDTA. Unincorporated nucleotides were removed with ProbeQuant G-50 columns per manufacturer’s instructions. 100ng of labeled probe DNA was run on a 1.5% agarose gel following each nick/translation reaction to ensure the majority of probe fragments were in the 300-800bp range. Up to 200ng total probe per cover slip was combined with 10ug salmon sperm DNA (ThermoFisher 15632011), 4ug human Cot1 DNA (ThermoFisher 15279011), 1/10 volume of 3M sodium acetate pH 5.2, and 2.5 volumes of 100% ethanol. Probes were allowed to precipitate for 30 minutes at −20°C, were centrifuged for 20 minutes at 4°C, maximum speed, washed twice with 70% ethanol and once with 100% ethanol, air dried, and resuspended in 6uL 100% formamide at 56°C. 6uL of 2x hybridization buffer (40% dextran sulfate in 8x SSC (20x SSC: 3M NaCl, 0.3M Sodium citrate)) was then added to each probe. Probes were denatured for 5 minutes at 80 degrees and snap-cooled on ice. Probes were then added to cover slips and denatured and hybridized to genomic DNA as noted above. The *CXCL* locus FISH probe utilized BAC RP11-243E9, the heterochromatic control probe utilized BAC RP11-134J16.

Imaging of FISH samples was performed at the Waitt Biophotonics Center at the Salk Institute. FISH samples were imaged on Zeiss Airyscan 880 microscopes using the Airyscan Fast mode (Huff 2016) at a resolution of 40nm in the x and y axes. Z sections were imaged every 160nm. Quantification of FISH data was performed using TANGO (Ollion et al. 2013) for FIJI (Schindelin et al. 2012). Nuclei and spot detection were performed with built in tools in TANGO. Image metrics analyzed in TANGO include: “Eroded Volume Fraction” and “Signal Quantification Layer” in Fig. 4, and “Distances” in Supplemental Fig. S3. Metrics were exported from TANGO as text files and statistical analysis and figure generation were performed in R using built-in tools (R Core Team).

### HiC analysis

Raw HiC library read alignment to human genome build hg38, valid read pair filtering, matrix assembly at various resolutions, and ICE normalization of said matrices were performed using HiC-pro with default settings (Servant et al. 2015). Biological replicates were pooled following valid read pair filtering, and pooled data sets were used for analysis except where noted.

For all direct comparisons of HiC data (topological domain boundary location comparisons, insulation scores, plotted contact matrices, log_2_ differential matrices) ICE normalized sparse matrix files were created containing only the subset of interacting bins that recorded reads in all data sets being compared. Read numbers at these bins were then quantile normalized in R using the normalize.quantiles() function in the preprocessCore package (Bolstad), allowing direct comparison of chromatin interactions between libraries with different read distributions and sequencing depths (Hsu et al. 2017).

Topological domain boundaries were called on normalized HiC data at 40kb resolution using the domain calling software published in Dixon et al. (Dixon et al., 2012).

HiC-Pro defined valid read pairs were used in conjunction with HOMER (Heinz et al. 2010) to run principal component analysis (PCA, *runHiCpca.pl -res 10000*), generate distance vs. interaction frequency plots (*makeTagDirectory*), define compartment boundaries (*findHiCCompartments.pl*), determine interaction correlations (*getHiCcorrDiff.pl -res 40000 - superRes 40000*), define distance-normalized chromatin interactions (*analyzeHiC -res 20000 - superRes 40000 -minDist 100000*), and to generate whole chromosome pairing plots (*analyzeHiC -res 400000000*).

CTCF anchored-type loops were called using HICCUPS (Rao et al. 2014).

Insulation scores were determined as follows: The genome was divided into 40kb segments. Insulation scores for each segment were defined as the number of normalized (ICE and quantile, see above) valid read pairs within a 500kb window centered on the segment of interest whose ends map to opposite sides of the segment of interest divided by the total number of valid read pairs whose ends both map within the 500kb window.

ΔPC1 domains were identified as follows: PCA was run at 10,000 base pair resolution on pooled HiC data using the *runHiCpca.pl* command in HOMER with the following settings: *-res 10000 -superRes 10000 -genome hg38*. Visual inspection showed that positive PC1 values corresponded to the gene-rich A compartment, and negative PC1 values corresponded to the gene-poor B compartment on all chromosomes and across all conditions. Genomic regions with PC1 score differentials between conditions greater than three standard deviations above the mean PC1 score differential between conditions were identified as potential ΔPC1 domains. PCA was then run on individual HiC biological replicates and only those potential ΔPC1 domains with a reproducible gain in PC1 value in each biological replicate were retained. Finally, reproducible ΔPC1 domains within 100kb of each other were merged into single continuous ΔPC1 domains which were used for downstream analysis.

### ChIP-seq analysis

Raw fastq files were aligned to the human genome build hg38 using bowtie (Langmead et al.) with the following parameters: *-m1 --best --strata*. Downstream processing of ChIP-seq data was performed using HOMER, except where noted. Uniquely mapped reads from high quality biological replicates were pooled for downstream analysis (Landt et al. 2012). Sequencing data was reorganized as a HOMER-formatted tag directory for each replicate and multiple reads mapping to the same base pair were collapsed to a single read using the *makeTagDirectory* command in HOMER with the following parameters: *-tbp 1*. ChIP peaks were called using the *findPeaks* command in HOMER with default parameters. Genes at ChIP peaks were identified using *annotatePeaks.pl* in HOMER, and the GenomicRanges package (Lawrence et al. 2013) in R.

SMC3-amassed enhancers were defined as follows: Enhancers were defined as H3K27ac peaks called as above. In order to identify enhancers with activation-dependent cohesin recruitment (SMC3-amassed enhancers), total unique SMC3 ChIP-seq reads mapping to enhancers were calculated using *annotatePeaks.pl* in HOMER. To directly compare binding strength between conditions read numbers at enhancers were quantile normalized across conditions using the preprocessCore R package. Those reads with a log2(normalized activated/normalized unstimulated read numbers) value greater than 1.5 were defined as SMC3-amassed. The GenomicRanges package in R was used to identify genes in contact with SMC3-amassed enhancers in conjunction with HOMER-defined chromatin interactions (detailed below). Enhancer-gene pairs were called as interacting if the center of one interaction anchor was within 10kb of an enhancer and the center of the other interaction anchor was within 50kb of a gene promoter.

### RNA-seq analysis

RNA-seq data was analyzed using the Tuxedo tools, except where noted. Raw fastq files were aligned to the human genome build hg38 using tophat2 (Kim et al. 2013) with the following parameters: *--library-type fr-firststrand -a 15*. Duplicated reads were removed using Picard tools command MarkDuplicates REMOVE_DUPLICATES=T, and RNA-seq quality metrics were assessed using Picard tools command CollectRnaSeqMetrics (http://broadinstitute.github.io/picard). Gene expression values were computed for each replicate across each condition using cuffdiff with an hg38 refflat file as reference with the following parameters: *--library-type fr-firststrand*. Subsequent analysis of gene expression and integration of gene expression data with other data types was performed in R.

### Metascape analysis (http://metascape.org)

Genes associated with various genomic features were identified using the GenomicRanges package in R and were analyzed for functional enrichment in the Metascape web portal using “Express Analysis” on default settings (Tripathi et al. 2015). Metascape gene set enrichment visualizations were performed in R.

### Bisulfite-seq analysis

Bisulfite converted DNA sequencing data was processed using the BSseeker2 software suite (Guo et al. 2013). A bisulfite-sequencing amenable hg38 reference genome was built using the *bs_seeker2-build.py* command. DNA sequences were aligned to the hg38 bisulfite sequencing-amenable genome build using the *bs_seeker2-align.py* command with the following options: *-m 6 -I 0 -X 800*. Cytosine methylation levels were determined using *bs_seeker2-call_methylation.py* with default settings. Awk was used to convert CGmap files to HOMER-compatible allC formatted files. HOMER-formatted tag directories were built using HOMER’s *makeTagDirectories* command with the following options: *-format allC -minCounts 0 -genome hg38*. Due to sequencing coverage-induced biases in DNA methylation meta-analysis (data not shown), awk was used to create HOMER-formatted tag directories containing only those cytosine residues covered by both unstimulated and PMA-activated neutrophil data sets. HOMER’s *annotatePeaks.pl* command was used with the *-ratio* option to determine DNA methylation levels at particular genomic features.

### Data visualization

Normalized HiC contact matrices presented in this paper were generated using HiCPlotter (Akdemir and Chin 2015). HiC interactions and ChIP-seq data in Fig. 4, Supplemental Figs. S4, and S6 were visualized using Sushi (Phanstiel et al. 2014). The remainder of linear genomic data was visualized using the Integrated Genomics Viewer (Robinson et al. 2011; Thorvaldsdóttir et al. 2013). FISH images were processed in FIJI. All other data was visualized using R.

### Data availability

Data sets generated in this study are available as a series in the GEO database under accession number GSE126758.

## Acknowledgements

We thank Alex Bortnick and other members of the Murre lab for editing the manuscript, and Yolanda Markaki and Kathrin Plath for assistance with the FISH protocol. This study was supported by funding from the CCBB (UL1TRR001442), the CIRM (RB5-07025), and the NIH to C.M. (U54DK24230, AI082850, AI00880 and AI09599) and V.N. (1 U01 AI124316), from the Frontiers of Innovation Scholars Program to MD, and by the Waitt Advanced Biophotonics Core Facility of the Salk Institute with funding from NIH-NCI CCSG: P30 014195 and the Waitt Foundation. TI was supported by the Uehara Memorial Foundation. High throughput sequencing was performed at the IGM Genomics Center, University of California, San Diego, La Jolla, CA.

## Author contributions

M.D. performed the majority of the experiments and analysis. Y.Z. processed HiC samples. A.H., H.L., T.I., and S.D. provided technical support and advice. M.D. and C.M. wrote the manuscript. V.N. and C.M. supervised the study.

