## Supplemental figures and legends for "Upon microbial challenge human neutrophils undergo rapid changes in nuclear architecture to orchestrate an immediate inflammatory gene program"

Denholtz Supplemental Figure S1

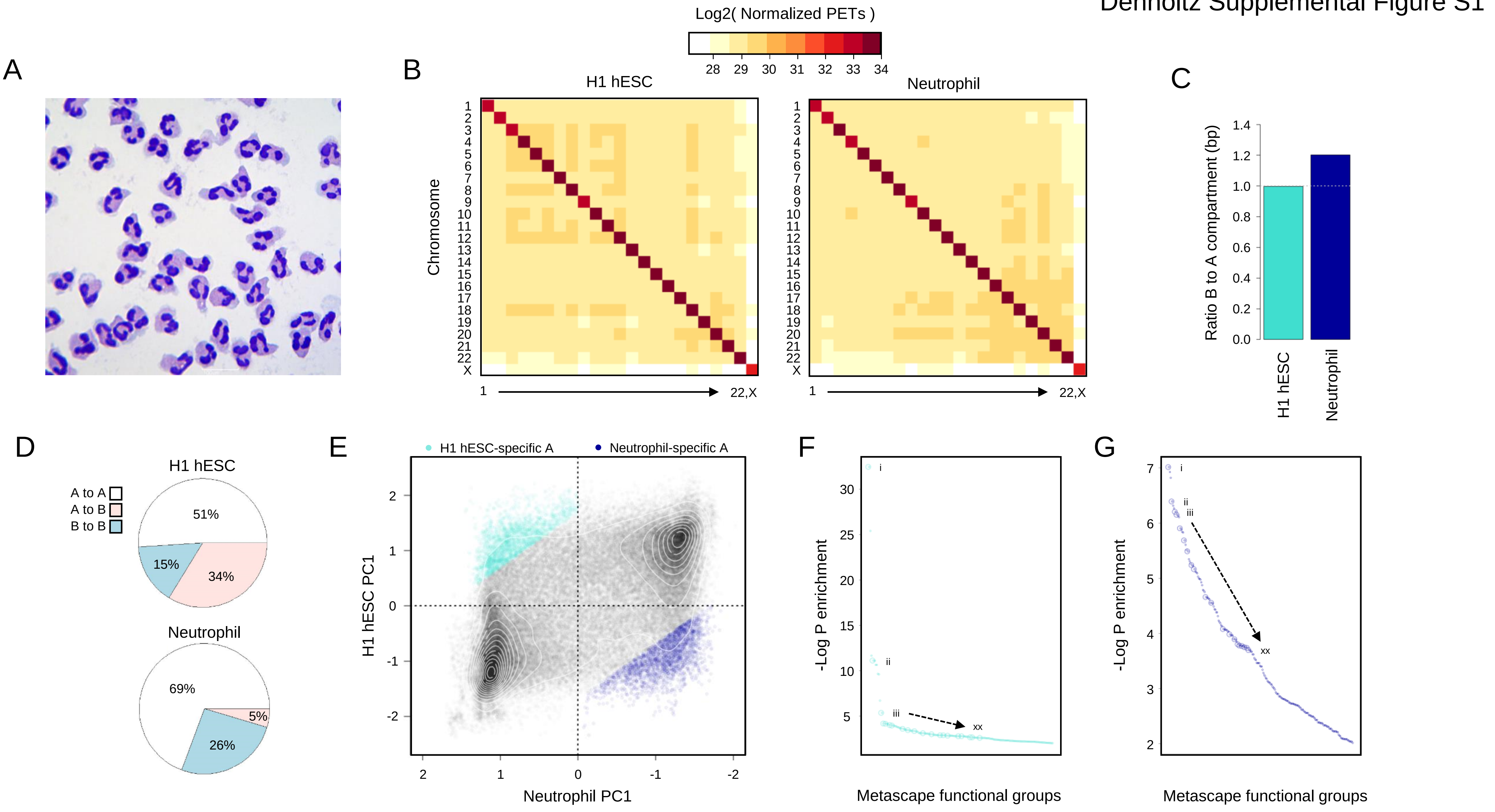

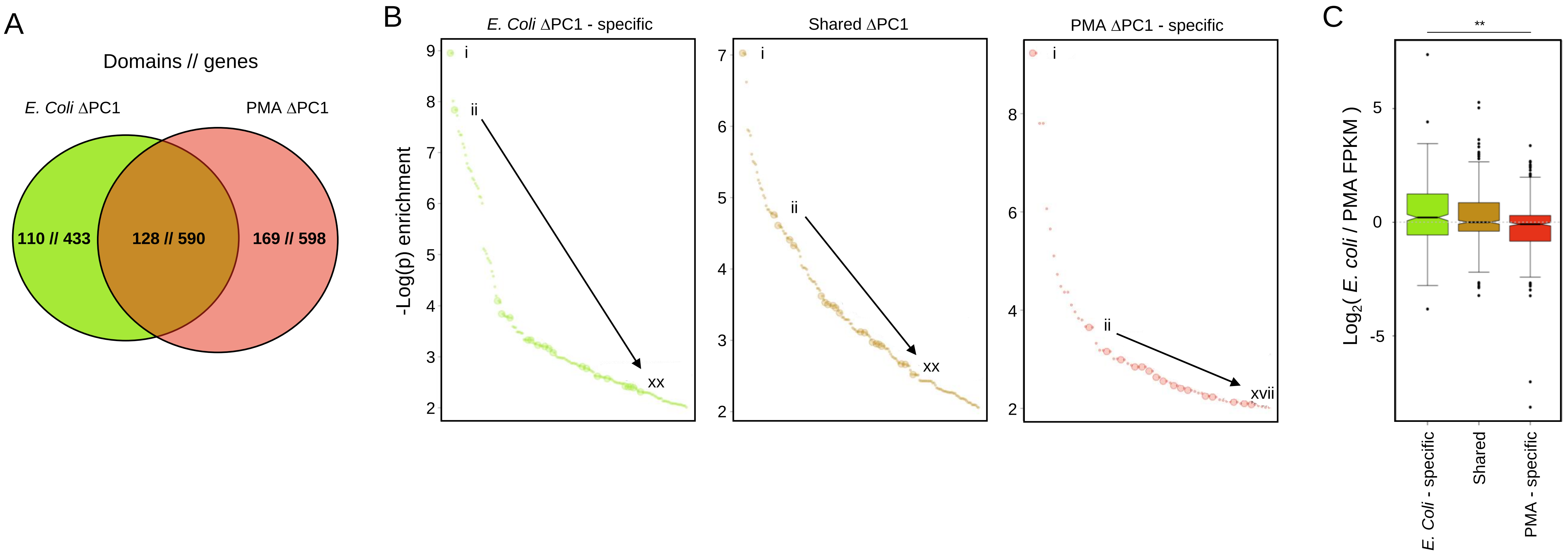
A
Domains // genes
E. Coli ΔPC1
PMA ΔPC1
110 // 433
128 // 590
169 // 598
B
E. Coli ΔPC1 - specific
Shared ΔPC1
PMA ΔPC1 - specific
-Log(p) enrichment
i
ii
xx
xvii
C
**
Log2( E. coli / PMA FPKM )
5
0
-5
E. Coli - specific
Shared
PMA - specific

A

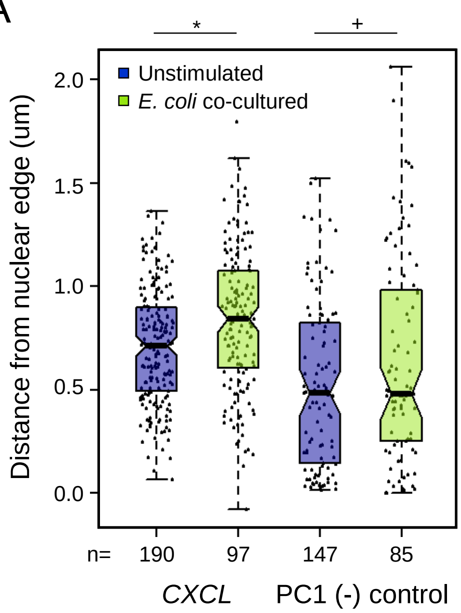
*
+
Unstimulated
E. coli co-cultured
Distance from nuclear edge (um)
2.0
1.5
1.0
0.5
0.0
n=
190
97
147
85
CXCL
PC1 (-) control

Denholtz Supplemental Figure S4

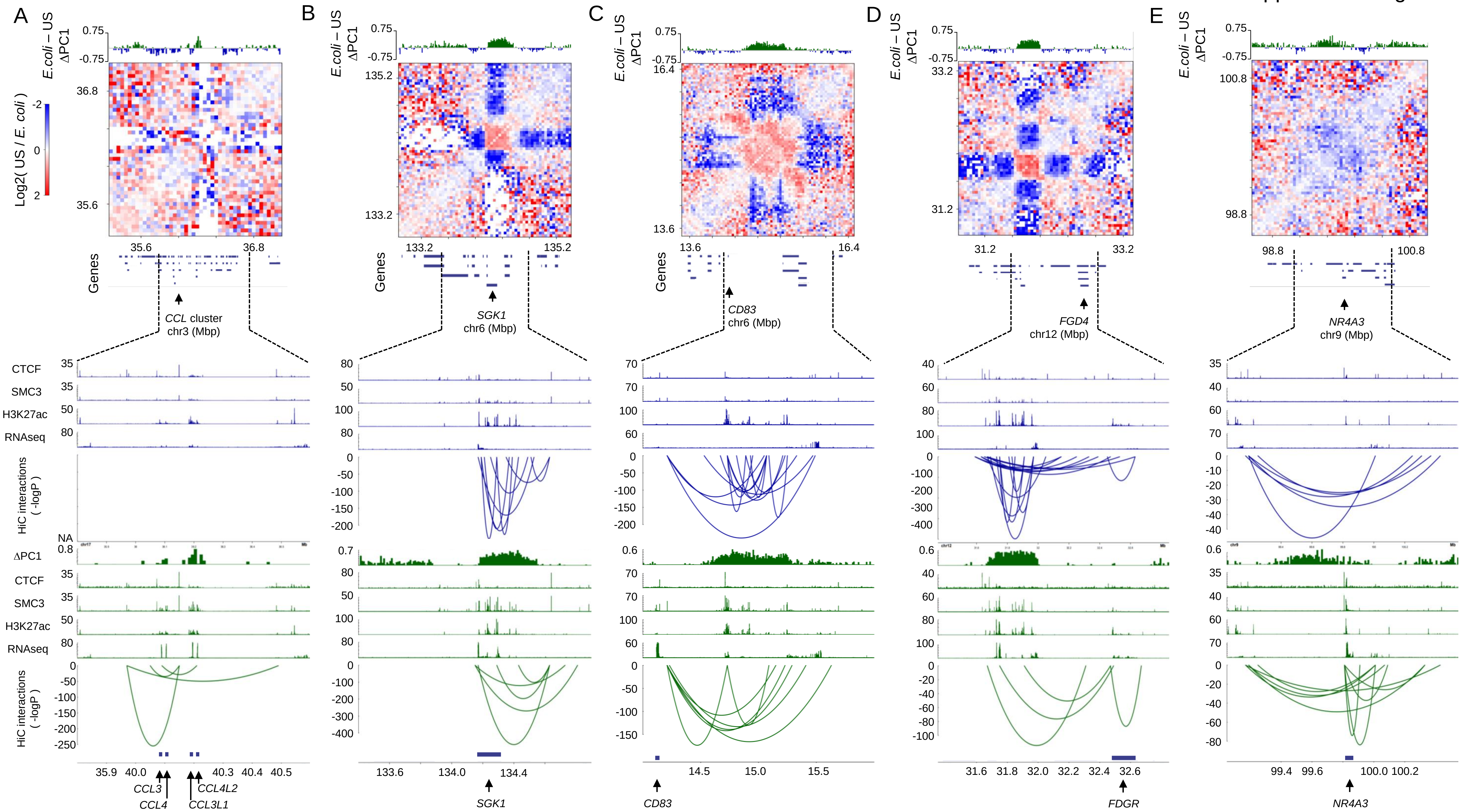

Denholtz Supplemental Figure S5

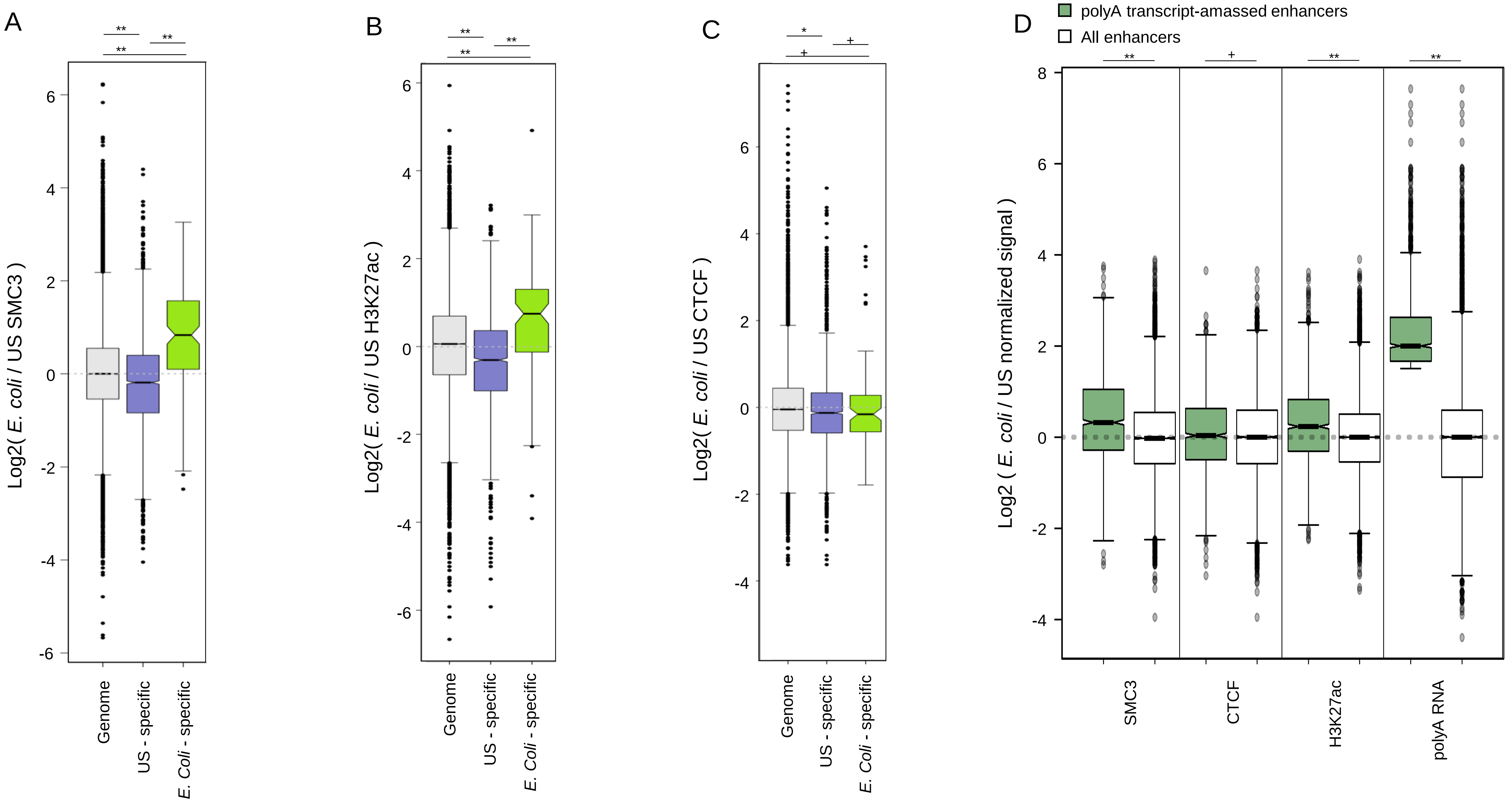

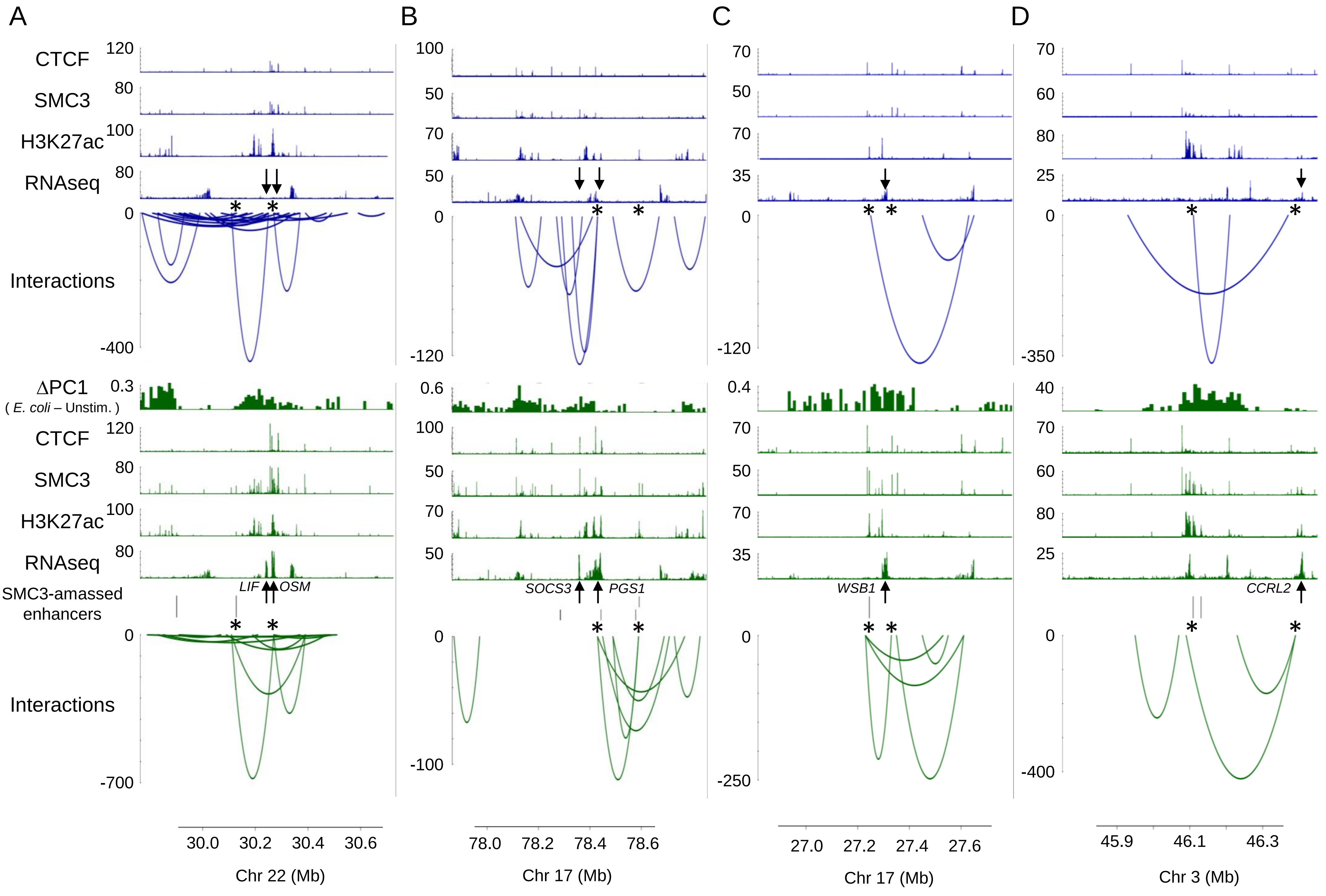
A
B
C
D
CTCF
SMC3
H3K27ac
RNAseq
Interactions
ΔPC1
( E. coli – Unstim. )
SMC3-amassed enhancers
LIF
OSM
SOCS3
PGS1
WSB1
CCRL2
Chr 22 (Mb)
Chr 17 (Mb)
Chr 17 (Mb)
Chr 3 (Mb)
30.0
30.2
30.4
30.6
78.0
78.2
78.4
78.6
27.0
27.2
27.4
27.6
45.9
46.1
46.3

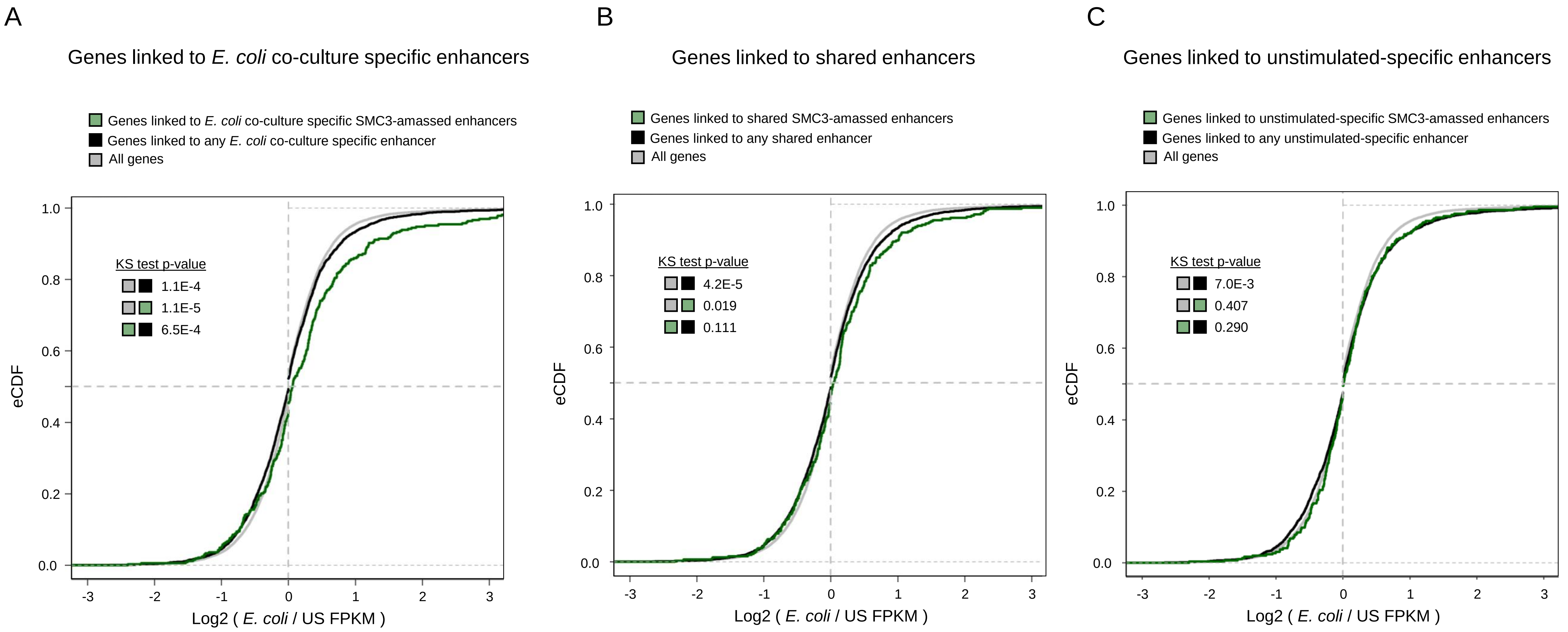
A
Genes linked to E. coli co-culture specific enhancers
Genes linked to E. coli co-culture specific SMC3-amassed enhancers
Genes linked to any E. coli co-culture specific enhancer
All genes
KS test p-value
1.1E-4
1.1E-5
6.5E-4
eCDF
Log2 ( E. coli / US FPKM )
B
Genes linked to shared enhancers
Genes linked to shared SMC3-amassed enhancers
Genes linked to any shared enhancer
All genes
KS test p-value
4.2E-5
0.019
0.111
eCDF
Log2 ( E. coli / US FPKM )
C
Genes linked to unstimulated-specific enhancers
Genes linked to unstimulated-specific SMC3-amassed enhancers
Genes linked to any unstimulated-specific enhancer
All genes
KS test p-value
7.0E-3
0.407
0.290
eCDF
Log2 ( E. coli / US FPKM )

**Supplemental Figure S1.** Changes in chromatin organization upon neutrophil differentiation.

(A) Wright-Giemsa stain of primary human neutrophils isolated from peripheral blood.

(B) Read normalized HiC matrices, each pixel represents normalized reads between chromosomes for H1 hESC on left and unstimluated human neutrophils on right.

(C) Ratio of total base pairs in the B compartment and total base pairs in the A compartment for H1 hESC (teal) and human neutrophils neutrophils (blue).

(D) Significant interchromosomal interactions broken down by compartment membership of anchor fragments for H1 hESCs (top) and human neutrophils (bottom).

(E) Scatterplot comparing PC1 values of H1 hESCs and neutrophils at 40kb resolution with density contour plot overlaid in white. Regions that flip from the B to A compartments and have PC1 differences greater than two standard deviations of the genome-wide mean PC1 difference are colored by cell type: teal = H1 hESC; blue = neutrophil.

(F) Metascape gene functional analysis for genes in H1 hESC-specific PC1 positive regions highlighted in E. See supplemental table 2 for gene functional group designations.

(G) Metascape gene functional analysis for genes in neutrophil-specific PC1 positive regions highlighted in E. See supplemental table 2 for gene functional group designations.

**Supplemental Figure S2.** Stimulation-specific ΔPC1 domains are independently regulated and contain distinct genes sets**.**

(A) Venn diagram illustrating the overlap between *E. coli* ΔPC1 and PMA ΔPC1 domains and their respective genes defined during *E. coli* co-culture and PMA activation. Genes within 100kb of ΔPC1 domains were considered.

(B) Metascape analysis for gene sets defined in A. Summary groups are marked by enlarged points, complete metascape results can be found in Supplemental table 2.

(C) Log2( *E.coli* co-culture / PMA-activated neutrophil FPKM ) gene expression values for genes sets defined in A. ** Wilcoxon rank-sum test p-value: <1E-10

**Supplemental Figure S3.** *CXCL8* and PC1 (-) control dual color FISH**.**

(A) FISH results showing the distance of the *CXCL8* locus and a control PC1 (-) FISH probe from the nuclear edge in unstimulated and *E. coli* co-cultured neutrophils. Wilcoxon rank sum test p-values: *=0.0002; +=0.2714.

**Supplemental Figure S4.** *E. coli* ΔPC1 domains lose dense intra-subdomain chromatin contacts in favor of long-range regulatory Interactions**.**

(A) *E. coli* ΔPC1 domain surrounding the *CCL* cluster. From top to bottom: PC1 differential (*E. coli* co-cultured – unstimulated PC1); $Log_2$(*E. coli* co-cultured/unstimulated) differential HiC contact matrix; protein coding genes; unstimulated CTCF, SMC3, and H3K27ac ChIP-seq, RNA-seq, and chromatin interactions; PC1 differential (*E. coli* co-cultured – unstimulated); *E. coli* co-cultured CTCF, SMC3, and H3K27ac ChIP-seq, RNAseq, and chromatin interactions. Only chromatin interactions with a logP value of <= -50 are shown.

(B) As in A for the *SGK1* locus.

(C) As in A for the *CD83* locus.

(D) As in A for the *FDG4* locus.

(E) As in A for the NR4A3 locus.

**Supplemental Figure S5.** Cell type-specific chromatin loops are associated with cell type-specific SMC3 and H3K27ac**.**

(A) $Log_2$( normalized *E. coli* co-cultured / unstimulated ) SMC3 ChIP-seq signal at all SMC3 peaks genome wide, at SMC3 peaks at unstimulated neutrophil-specific interaction anchors, and at SMC3 peaks at *E. coli* co-culture-specific interaction anchors.

(B) As in A, for H3K27ac ChIP-seq peaks and ChIP-seq signals.

(C) As in A, for CTCF ChIP-seq peaks and ChIP-seq signals.

(D) $Log_2$ ratio (*E. coli* co-cultured / unstimulated) of normalized ChIP-seq and RNA-seq signals at enhancers that amass polyadenylated transcripts and all enhancers genome-wide.

Wilcoxon rank sum test: **  <2E-16; * <1E-4; x = 2E-3; + not significant

**Supplemental Figure S6.** SMC3-amassed enhancers form de novo chromatin loops with inflammatory gene promoters**.**

(A) Example of *E. coli* co-culture-induced SMC3 binding at a pre-existing enhancer resulting in increased interaction strength between the enhancer and the *LIF* and *OSM* gene locus, resulting in their transcriptional up-regulation. CTCF, SMC3, and H3K27ac ChIP-seq, and RNAseq signals, chromatin interactions, and changes in PC1 scores are shown for unstimulated neutrophils in blue, and *E. coli* co-cultured neutrophils in green. Arrows show example genes, asterixis show position of loop anchors connecting SMC3-amassed enhancers to their target genes in *E. coli* co-cultured neutrophils.

(B) As in A, for the *SCOS3*/*PGS1* locus, illustrating de novo loop formation between an SMC3-amassed enhancer cluster and their target genes.

(C) As in B, for the *WSB1* gene locus.

(D) As in B for the *CCRL2* gene locus.

**Supplemental Figure S7.** Pre-existing and *de novo* formed enhancers both rely on cohesin recruitment to activate gene expression.

(A) $Log_2$ differential (*E. coli* co-cultured / unstimulated FPKM) gene expression cumulative distribution frequency plots for all genes, genes linked to any H3K27ac-defined enhancer specific to *E. coli* co-cultured neutrophils, and for genes linked to SMC3-amassed H3K27ac-defined enhancers specific to *E. coli* co-cultured neutrophils.

(B) As in A, for enhancers shared between unstimulated and *E. coli* co-cultured neutrophils.

(C) As in A, for enhancers identified only in unstimulated neutrophils.
